# AMPA receptor desensitization shapes gain control in the lateral geniculate nucleus

**DOI:** 10.1101/2024.03.12.584576

**Authors:** Sonia Ruggieri, Tim Gollisch, Jakob von Engelhardt

**Affiliations:** Institute of Pathophysiology, University Medical Center, Johannes Gutenberg University Mainz, Mainz, Germany; Department of Ophthalmology, University Medical Center Göttingen, Göttingen, Germany

## Abstract

Synaptic short-term plasticity and EPSP summation determine the processing of visual information by dLGN relay cells. *In vitro* studies suggested that desensitization of AMPA receptor results in pronounced short-term depression in retinogeniculate synapses thereby influencing activation of relay cells by retinal ganglion cell spikes. However, whether retinogeniculate synapses depress and whether AMPA receptors desensitize *in vivo* is controversial. We here tested the role of AMPA receptor desensitization in processing of visual information by generating a computational model of a dLGN relay cell containing AMPA receptors with different levels of desensitization. In addition, we investigated the *in vivo* activity of dLGN relay cells in wildtype and CKAMP44^−/−^ mice. CKAMP44 is an auxiliary subunit that increases desensitization of AMPA receptors in retinogeniculate synapses. The comparison of responses of relay cells in the two genotypes therefore allowed us to obtain information about the contribution of AMPA receptor desensitization in processing of visual information. The simulations and the *in vivo* data show that AMPA receptor desensitization affects relay cell activity by decreasing input gain and response gain. In addition, desensitization sharpens the response of relay neurons to moving gratings in the temporal domain by decreasing the firing rate when the non-preferred stimulus is presented. Altogether, we uncover a complex role of CKAMP44 and AMPA receptor desensitization in information processing of dLGN relay neurons *in vivo*.

## Introduction

The input that a neuron receives is usually not simply relayed to the next neuron but filtered depending on dynamic changes in synaptic strength as well as passive and active membrane properties. Summation of several EPSPs is usually required to elicit action potentials and saturation limits the maximal firing frequency. Synaptic strength on the other hand changes dynamically with input frequency. An interplay of summation, saturation and synaptic short-term plasticity (STP) therefore determines signal transmission and filter properties of neurons.

Relay neurons of the dLGN and their main driver input, i.e. the retinogeniculate synapse, have been particularly valuable as a model system in which the variables that determine signal transmission were studied in detail^1–9^. dLGN relay cells (RCs) receive 1-2 dominant inputs from retinal ganglion cells (RGCs) with a high current amplitude of >1200pA resulting from a large number of release sites^4^ and high release probability^5^. AMPAR desensitization and high release probability contribute to the pronounced short-term depression (STD) that was observed *in vitro*^3,4,10,11^. However, whether or not retinogeniculate synapses depress *in vivo* is controversial^6,12,13^. A likely explanation for a lower depression of synapses *in vivo* than *in vitro* is a lower vesicle release probability^1^. This also questions the contribution of AMPAR desensitization in signal transmission as the role of desensitization in synaptic short-term plasticity decreases with the vesicle release probability^5^.

The role of EPSP summation in signal processing of dLGN neurons, on the other hand, is more evident. Spike transmission rate is significantly higher for the second of two action potentials due to EPSP summation if retinogeniculate synapses are activated at high frequency (>30 Hz)^6–8^. This filtering is highly relevant as the second spike carries more visual information than the first^14,15^. Importantly, EPSP summation and synaptic depression are not mutually exclusive but act together to fine-tune signal transmission. Summation ensures efficient activation of dLGN neurons by weak visual stimuli (i.e. when a retinal ganglion cell fires two action potentials) but not by individual spikes that carry little information (i.e. noise). Summation bears on the other hand the risk that dLGN neurons saturate during high retinal ganglion cell activity. Saturation may be prevented by AMPAR desensitization-mediated STD in retinogeniculate synapses.

We recently showed that the AMPAR auxiliary subunit CKAMP44 increases desensitization and reduces the rate of recovery from desensitization^16^. Retinogeniculate synapses display facilitation instead of depression in CKAMP44 knockout (CKAMP44^−/−^) mice confirming the relevance of desensitization in short-term plasticity *in vitro*. Moreover, deletion of CKAMP44 increased the responsiveness of dLGN neurons which can be explained by a reduced AMPAR desensitization and fast recovery from desensitization. Considering that CKAMP44 alters filter properties of retinogeniculate synapses *in vitro*, we hypothesize that AMPAR desensitization exerts a more complex influence on signal transmission than simply decreasing the responsiveness of dLGN relay neurons.

We here tested the role of AMPAR desensitization in signal transmission in retinogeniculate synapses. To this end, we developed a detailed computational model of dLGN neurons with retinogeniculate synapse containing AMPARs with gating kinetics observed in wildtype or CKAMP44^−/−^ mice^16^, i.e. AMPAR with strong and weak desensitization, respectively. Results from simulations allowed to scrutinize key factors involved in retinogeniculate synapse function such as glutamate spillover. Moreover, the simulations suggested that reduced AMPAR desensitization has a complex influence on response tuning curves of dLGN relay neurons by decreasing input and response gain. To confirm the relevance of AMPAR desensitization for signal transmission, we performed *in vivo* tetrode recordings in awake head-fixed adult wildtype and CKAMP44^−/−^ mice while presenting a battery of visual stimuli. Absence of CKAMP44 indeed altered contrast response curves and drifting grating tuning curves consistent with altered filter properties of dLGN neurons corroborating findings from simulations. Our results demonstrate an unexpected complex and strong influence of AMPAR desensitization on signal transmission in the visual system.

### AMPARs desensitization increases STD in a computational model of a retinogeniculate synapse

To better understand the role of AMPAR desensitization in filtering of visual information in the thalamus, we developed a detailed computational model of a dLGN neuron. We used AMPAR-mediated currents that we had previously recorded from patches of dLGN RCs to derive two kinetic models of AMPARs with different levels of desensitization, i.e. 1) currents recorded from wildtype mice for a model with pronounced desensitization and slow recovery from desensitization and 2) currents recorded from CKAMP44^−/−^ mice for a model with little desensitization and fast recovery from desensitization^16^. Two sets of rate constants of a 9-states AMPARs kinetic model^5,17^ were determined (**Fig. 1A**), which allowed us to simulate currents that resembled those recorded from patches of dLGN neurons of wildtype and CKAMP44^−/−^ mice (**Fig. 1B-F**). Gating kinetics of simulated currents were similar to experimentally determined gating kinetics (**Fig. 1G-H**).

**Figure 1-.**
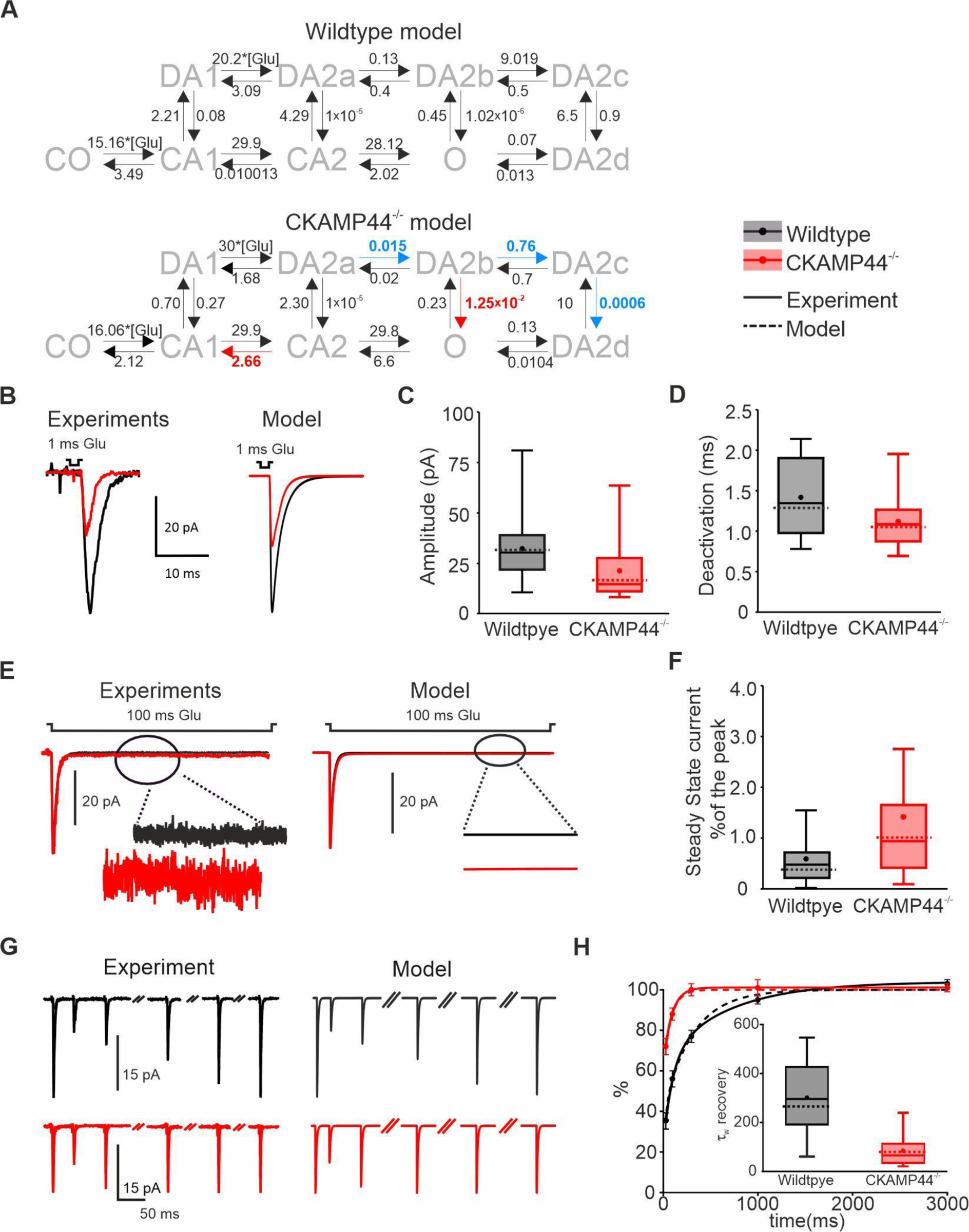
Kinetic models for AMPAR-mediated currents of wildtype and CKAMP44^−/−^ mice. **A)** *K*inetic model*s with* C0, CA1 and CA2 indicating conformational closed states, DA1 and DA2a-d desensitized states and O open state. Glutamate concentration is indicated with [Glu]. Values indicate the rate constants of each conformational transition of the receptor. Rate constants that increase or decrease more than one order of magnitude in the CKAMP44^−/−^ *receptor* model are highlighted in red. **B)** Currents in response to 1mM glutamate applied for 1ms. Experimental traces on the left, model data on the right. Peak amplitude **(C)** and deactivation time constant **(D)** of currents from recordings of *Chen et al.*^16^. The dashed line shows the value of the computational model. **E)** Currents in response to 1mM glutamate applied for 100ms. Experimental traces on the left, model data on the right. **F)** Quantification of steady state amplitude as a percentage of peak amplitude of recorded currents. The dashed line shows the value of the computational model. In all plots, the dot represents the experimental mean value. **G)** Currents in response to two 1ms pulses of 1mM glutamate. Experimental traces on the left, simulated data on the right. **H)** Quantification of the recovery from desensitization for both experimental and simulated data. The inset shows the time constant of the recovery from desensitization (τ_w_ recovery). Experimental data: median ± IQR, dot = mean. Dashed line = result from the model.

The analysis of all conformational states in response to two 1mM glutamate pulses with an interpulse-interval of 30 ms obtained from the simulations showed that the presence of CKAMP44 increases in particular the probability of occupancy of desensitized states and consequently reduces the open probability of AMPARs during the second glutamate pulse (*Extended Data Fig. 1A-C*).

We used the kinetic models of AMPARs as well as previously described anatomical and functional data^5,16,18^ to develop a 3D computational model (NEURON program, Yale) of a dLGN neuron with a retinogeniculate synapse containing wildtype or CKAMP44-lacking AMPARs. The aim in the generation of the computational model was to analyze the role of AMPAR desensitization kinetics in retinogeniculate synapse function. In addition, simulations of dLGN neuron activity containing AMPA receptors with different levels of desensitization facilitates the interpretation of data from *in vivo* recordings of wildtype and CKAMP44^−/−^ mice. Absence of CKAMP44 reduces AMPAR-mediated current amplitude by 43% in retinogeniculate synapses consistent with the role of CKAMP44 in increasing synaptic AMPAR number^16,19,20^. To take the experimentally observed^16^ decrease in AMPAR-mediated current amplitude into account, we reduced the peak conductance in the model of retinogeniculate synapses lacking CKAMP44 by 43% when compared to the conductance in the wildtype model. Of note, the wildtype and CKAMP44^−/−^ retinogeniculate synapse models differed only in the AMPAR conductance and kinetics. All other anatomical and functional parameter were the same in the models of the two genotypes.

We then compared synaptic STP of the two models with the previously *in vitro* determined synaptic STP of retinogeniculate synapses in wildtype and CKAMP44^−/−^ mice^16^. To this end, we simulated current pairs by activating the model synapse twice with different inter-pulse intervals. STP of the models indeed resembled the experimentally determined STP as evidenced by the similarity of the simulated and recorded currents (**Fig. 2A**) as well as the quantification of the paired-pulse ratios (PPRs) of the currents (**Fig. 2B-C**). The simulations were performed with release probability (Pr= 0.4) that were lower than the previously reported Pr of 0.7 estimated from *in vitro* studies^5^. We used this lower Pr as it likely better reflects the *in vivo* Pr^1^. As expected, the difference in PPRs of the two models increased with Pr (**Extended data 3**), but there was a considerable difference in PPR also when performing simulations with lower Pr than 0.7 (**Extended data 3**).

**Figure 2.**
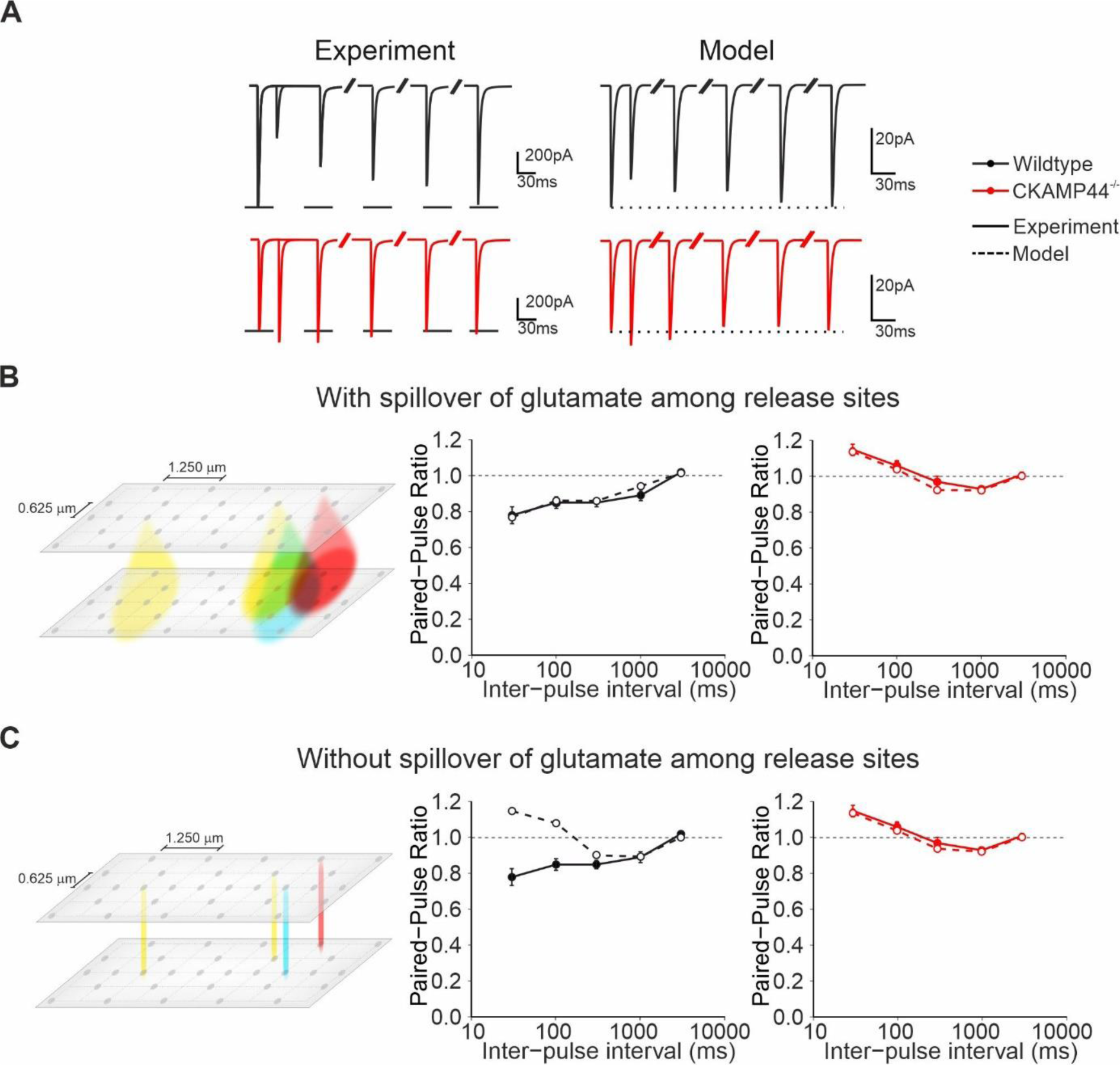
Synaptic short-term plasticity in computational models of relay cells. **A)** On the left example experimental traces of currents evoked by pairs of stimulations of retinogeniculate synapses with different interpulse-interval. On the right traces obtained from the retinogeniculate synapse model. Paired-pulse ratio of AMPAR-mediated currents with glutamate spillover **(B)** and without glutamate spillover **(C)**. Glutamate spillover is required in the model of wildtype synapses to simulate short-term plasticity observed in experiments. In contrast, spillover of glutamate does not significantly affect short-term plasticity in the model of synapses in CKAMP44^−/−^ mice. The continuous line represents the experimental data and the dashed line represents the data from the model. Experimental currents and paired-pulse ratio values (mean and SEM) are from Chen et al^16^.

Crosstalk between release sites may increase the relevance of AMPAR desensitization for STP. It has been suggested that the geometry of the giant retinogeniculate synapses prevents fast diffusion of glutamate outside the cleft such that glutamate can spillover from one release site to neighboring release sites. Glutamate receptors may therefore be desensitized not only in active but also in non-active release sites, which could depress current amplitude during subsequent activations. To analyze the relevance of glutamate spillover for STP in retinogeniculate synapses, we carried out the same simulations as above while setting glutamate spillover to zero. In this scenario, AMPARs are activated only by glutamate that is released from the opposing release site. Preventing glutamate spillover in the simulations increased PPRs considerably in the wildtype model as suggested previously by Budisantoso and colleagues^5^. Interestingly, the PPRs of the CKAMP44^−/−^ retinogeniculate model with and without glutamate spillover were almost similar. This suggests that glutamate spillover is only relevant for STP in retinogeniculate synapses containing CKAMP44-bound AMPARs, which recover slowly from desensitization.

### Desensitization of AMPARs reduces input and response gain of a computational model of a dLGN neuron

Investigation of firing probability in *in vitro* electrophysiological analyses showed that deletion of CKAMP44 increases the responsiveness of dLGN neurons. The frequency-dependence of this effect suggested that AMPAR desensitization-mediated STD affects filter properties of retinogeniculate synapses^16^. To investigate the role of AMPAR desensitization in signal transmission in more detail, we activated the RGC axon in the computational model with 7 stimuli at different frequencies and analyzed currents, open probability and desensitization of AMPARs as well as firing of the dLGN neuron (**Fig. 3 A-E**). As expected, synaptic STD was more pronounced in the wildtype than in the CKAMP44^−/−^ neuron model, which resulted from the more pronounced reduction in open probability and increase in desensitization state of CKAMP44-bound AMPARs. At low firing rates of the RGC axon (20-30 Hz), the wildtype dLGN neuron fired action potentials in contrast to the CKAMP44^−/−^ neuron model (**Fig. 3E**). This can be explained by the higher AMPAR number in the wildtype than the CKAMP44^−/−^ retinogeniculate synapse (note that the current amplitude at the first stimulus is higher in the wildtype than the CKAMP44^−/−^ neuron in **Fig. 3A**). At higher input frequencies, the CKAMP44^−/−^ model neuron displayed higher numbers of action potential than the wildtype model (**Fig. 3E**). Similar observations were made, when stimulating RGC axon with 5 or 10 stimuli instead of 7 (**Extended Figure 2**). AMPARs were fully activatable (i.e. not desensitized) before the stimulus train due to absence of pre-stimulus activity. The model may resemble in this respect the *in vitro* brain slice condition, in which activity levels are usually also very low such that AMPARs can fully recover from desensitization. However, the relatively high background (noise) activity of RGCs *in vivo* may result in a constant level of AMPAR desensitization^1,6^. To test the role of desensitization in signal transmission with *in vivo*-like background activity, we stochastically activated RGC axon in the model with 10 Hz for 1.5 s (background) prior to the higher frequency stimulus train. Background activity reduced AMPAR current amplitude (**Fig. 3F**) due to a reduced open probability (**Fig. 3G**) and increased number of AMPARs in the desensitized state (**Fig. 3H**). This effect was more pronounced in the wildtype model. Background activity induced in fact so strong desensitization of AMPARs such that the stimulus train did not elicit any action potential in the wildtype model neuron at any frequency (data not shown). Therefore, we increased the number of AMPARs in the retinogeniculate synapses of both models by a factor of 10%. The CKAMP44^−/−^ model neuron fired more action potentials than the wildtype neuron across all frequencies of RGC firing in the simulation with background activity (**Fig. 3J**). Hyperbolic ratio fitting of the input/output response function showed that C50 is lower and R_max_ higher in the CKAMP44^−/−^ model compared to the wildtype model (**Fig. 3J**). The reduction in the C50 and increase in R_max_ was also observed in simulations carried out with 5 and 10 pulses in the high frequency stimulus trains (**Extended Figure 2**). Therefore, based on the computational model, we predict that desensitization exerts a divisive control of input gain (reduction of C50) and response gain (increase in the maximal response amplitude)^21^.

**Figure 3.**
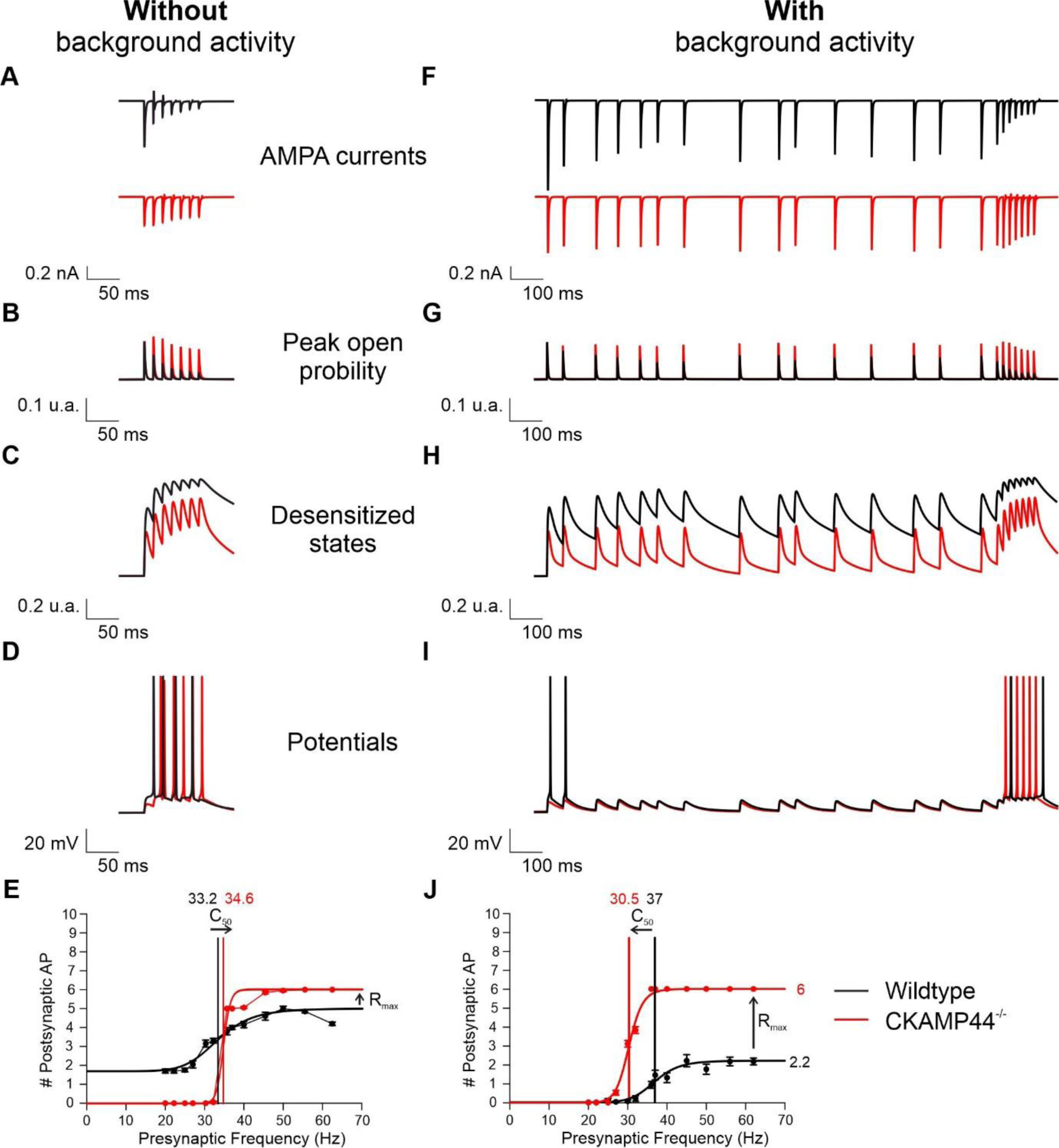
Input-output relationship in computational models of relay cells. **A)** AMPAR-mediated currents elicited by 7 stimuli at 50Hz. Sum of all AMPARs that during the stimulus are **(B)** desensitized and **(C)** open. **D)** EPSPs and action potentials elicited by the high frequency stimulus. **(E)** Input-output function of model neurons with wildtype and CKAMP44^−/−^-like AMPARs in retinogeniculate synapses. **(F-J)** same as **(A-E)** but in presence of 2s of background activity. Input-output curves in **(E** and **J)** were fitted by hyperbolic functions. ((E) R^2^_wt_=0.95, R^2^_CKAMP44_^−/−^=1.0, (J) R^2^_wt_=0.99, R^2^_CKAMP44_^−/−^=0.98).

### CKAMP44 does not influence receptive field shape of RCs *in vivo*

To test our prediction derived from the simulations about the role of AMPAR desensitization in processing of visual information, we performed tetrode recordings of single unit activity in the dLGN of awake head-fixed wildtype and CKAMP44^−/−^ mice (**Fig. 4A**). We presented a battery of different visual stimuli and recorded activity from ≈ 500 RCs in mice of each genotype. The complete data set of recorded units was then filtered depending on the visual stimulus considered (i.e. white noise, drifting gratings and contrast).

**Figure 4.**
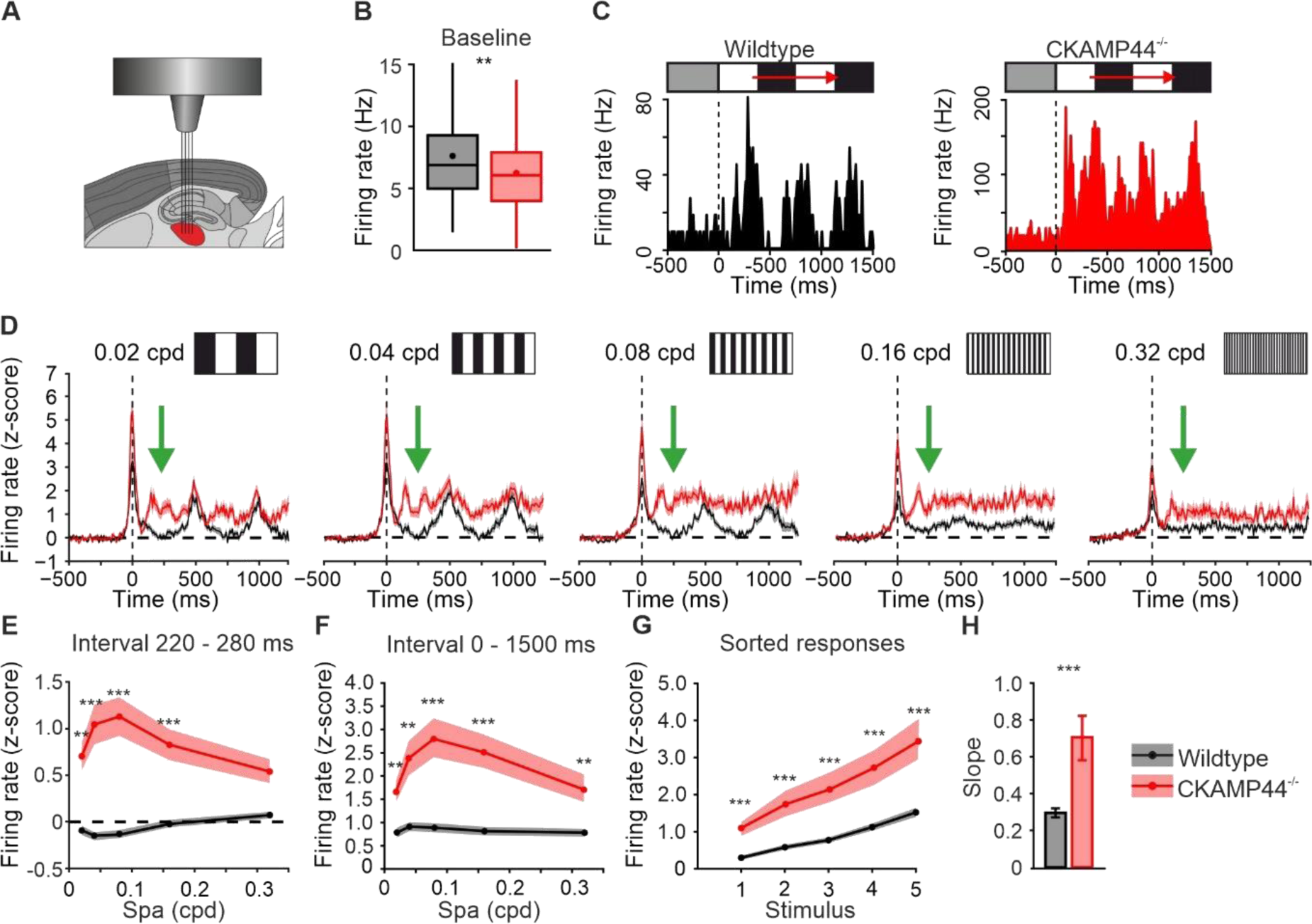
In vivo responses of relay cells to drifting gratings. **A)** Implantation of tetrodes in the dLGN (indicated in red). **B)** Background firing rate. Median ± IQR, dot = mean. Number of units: Wildtype: n=157, CKAMP44^−/−^: n=166. Wilcoxon test: W=15431.5; p=0.004. **C)** Activity of example units from wildtype (black) and CKAMP44^−\−^ mice in response to drifting gratings. **D)** Average of responses to drifting gratings at different spatial frequencies. The green arrow indicated where the valley amplitude was measured. **E)** Mean firing rate during the first valley (220-280 ms) at different spatial frequencies (Wilcoxon test: spa=0.02cpd W=10296, p=0.003; spa= 0.04cpd W=11016, p=0.00003; spa=0.08 cpd W=11541, p=0.0000005; spa=0.16 W=10494, p=0.0004, spa=0.32 W=8779, p=0.18). **F)** Mean firing rate during the entire stimulus period (0-1500 ms) at different spatial frequencies (Wilcoxon test: spa=0.02cpd W=6664, p=0.002; spa= 0.04cpd W=6831.5, p=0.005; spa=0.08 cpd W=6029, p=0.00004; spa=0.16 W=6045, p=0.00004, spa=0.32 W=6567.5, p=0.0012). **G)** Mean firing rates during the entire stimulus period (0-1500 ms) sorted for each unit from lowest to highest response (Wilcoxon test: stimulus 1 W=5868.5, p=0.000012; stimulus 2 W=6239, p=0.00016; stimulus 3 W=6529.5, p=0.0009; stimulus 4 W=6466, p=0.0006, stimulus 5 W=6364, p=0.0004). In panels E, F, and G, lines represent mean values and the shadows SEM. **H)** Bar plot of the slope from the linear fitting of the sorted spatial frequency tuning curves (Wilcoxon test: W=6284, p=0.0002, mean ± SEM). Number of units for all plots in the figure: Wildtype n=123, CKAMP44^−/−^ n=139 (*p < 0.05, **p < 0.01, ***p < 0.001).

To estimate whether deletion of CKAMP44 affects the structure of receptive fields of dLGN RCs, we analyzed responses to a spatiotemporal white-noise stimulus. Units that displayed clear receptive-fields were then selected for Spike Triggered Average (STA) analysis and classified as ON- and OFF-responsive units based on the polarity of their receptive field^22^ (*Extended data Fig. 4*). Spatial STA was then fitted by a 2-D Gaussian function and the radii were calculated as the standard deviations obtained from the Gaussian profiles along the x- and y-axis (*Extended data Fig. 4A-C*). Neither radius nor ratio of horizontal and vertical axis were altered in mice with CKAMP44 deletion, suggesting that size and the typical circular shape of the receptive field are preserved in CKAMP44^−/−^ units. Temporal STA was analyzed in a temporal window of 264ms preceding the spike of responsive RCs. There was no statistical difference between genotypes in the time course of stimulus amplitude as well as no change in the biphasic indices for ON-response and OFF-response centers (*Extended data Fig. 4F*). These findings suggest that desensitization of AMPARs in retinogeniculate synapses exerts little or no influence on encoding of receptive field structure in the dLGN. This may not be too surprising as previous studies showed that receptive fields are not transformed by RCs but are very similar to the receptive fields of their corresponding RGCs^23^. Moreover, unaltered receptive field structure is consistent with normal retinal function in CKAMP44^−/−^ mice. Indeed, we had previously shown that CKAMP44 exerts no influence on retinal function and AMPAR-mediated currents in RGCs^16^.

### CKAMP44 affects spatial frequency tuning curves of dLGN RCs *in vivo*

Our simulations of RC activity in response to RGC firing (**Fig. 3**) and our previous analysis of RC activity *in vitro*^16^ suggested that the magnitude of the influence of CKAMP44 (and thus AMPAR desensitization) depends on the activity rate of RGCs. To test this prediction, we investigated with in-vivo recordings spatial frequency tuning curves of dLGN RCs activity in response to drifting gratings (2Hz) with different spatial frequencies (0.02-0.32 cycles per degree; cpd). Of note, the mean firing rate of RCs in the 2s preceding the stimulus onset was slightly but significantly higher in wildtype than in CKAMP44^−/−^ mice (**Fig. 4B**).

For each responsive unit, we quantified the responses to drifting grating stimuli at their preferred orientation. When stimulating with moving gratings, the latency of the first peak of the responses of units shows considerably jitter (in particular due to the polarity and position of the receptive field of the unit) (**Fig. 4C**). To better investigate RCs response structure, we shifted activity profiles of all recorded units in time such that the first peak occurred at 0ms (*Extended data Fig. 5A*). There was a clear 2 Hz modulation of the average responses of units in wildtype (**Fig. 4D**). The 2 Hz modulation appeared to be less pronounced in the responses in CKAMP44^−/−^ mice (**Fig. 4D**). This observation was substantiated by the comparison of firing rates during the valleys of the responses. While firing rate was back to baseline firing rate during the valley in wildtype mice, there was a significantly increased firing rate during the valley in CKAMP44^−/−^ mice (**Fig. 4D-E**). Moreover, the lower 2 Hz modulation of the responses in CKAMP44^−/−^ mice is evidenced by a reduced ratio of the response amplitude of the second peak to the activity during the valley in CKAMP44^−/−^ mice when compared to wildtype, especially for low spatial frequencies of the drifting gratings. (*Extended data Fig. 5C)*. This result shows that CKAMP44 is relevant for encoding temporal information of visual stimuli presumably by allowing signal transmission during presentation of the preferred stimulus (e.g. On) but curtailing responses (and preventing activation by noise) during presentation of the non-preferred stimulus (e.g. Off) due to effective desensitization of AMPARs.

**Figure 5.**
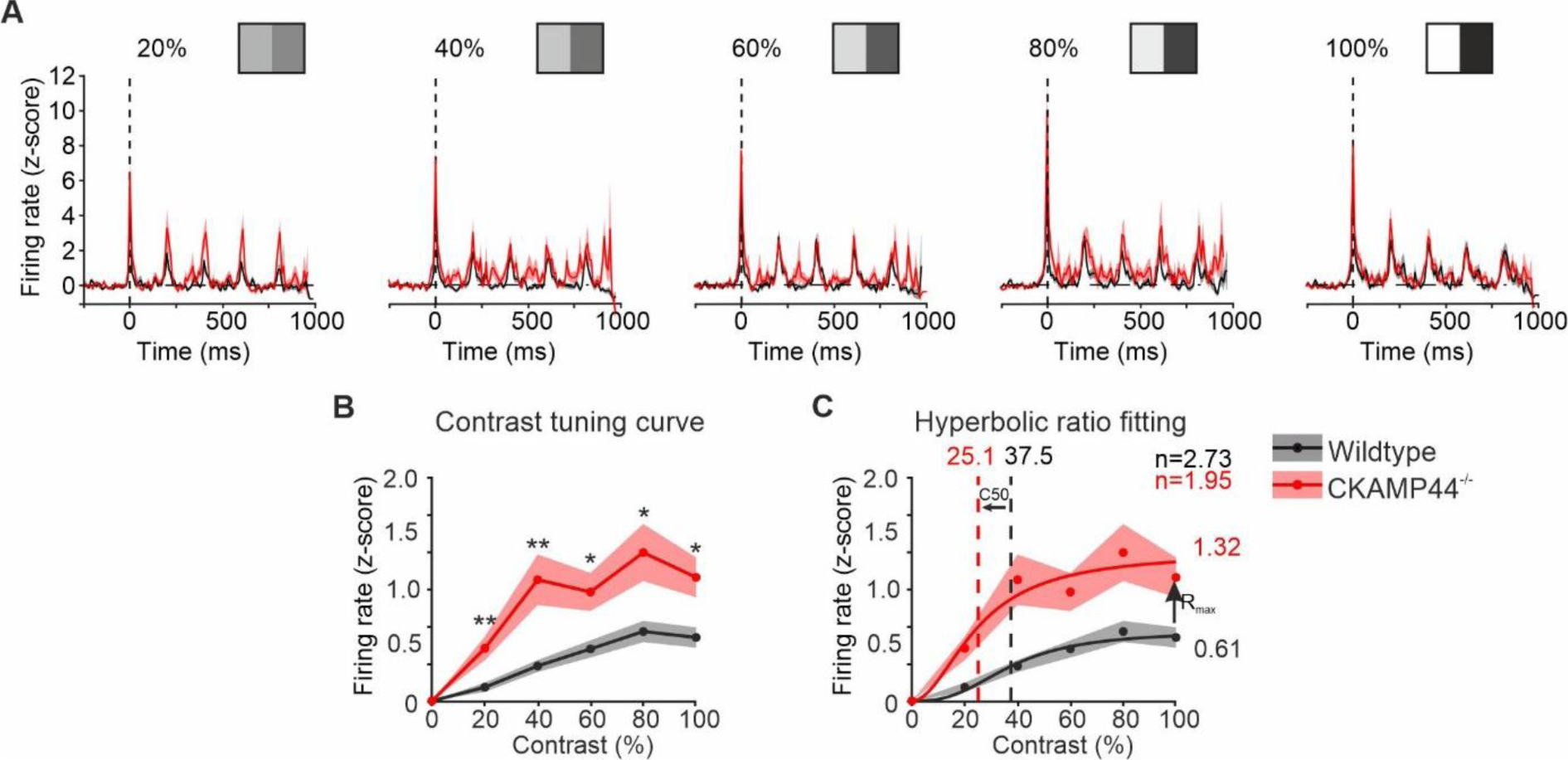
Contrast tuning curves of relay cells. **A)** Average unit activity at different contrast. The continuous line indicates the mean, whereas the shadow represents SEM. (n: wt=85, CKAMP44^−/−^=54). **B)** Contrast tuning curve with mean and SEM (Wilcoxon Test: Contr.20% - W=1302 p=0.008, Contr.40% - W=1210 p=0.0018, Contr.60% - W=1327 p=0.01, Contr.80% - W=1320 p=0.01, Contr.100% - W=1334 p=0.013). **C)** Hyperbolic fits to contrast tuning curves with C50 and R_max_ and n (goodness of the fit: R^2^ =0.98, R^2−/−^=0.97).

Deletion of CKAMP44 increased not only firing rate during the valley of the response, but also peak and mean firing rates (**Fig. 4F** and *Extended data Fig. 5C).* Previous results from *in vitro* experiments indicated that CKAMP44 strengthens low pass filter properties of retinogeniculate synapses^16^. This would suggest that CKAMP44 reduces signal transmission more when RGCs fire at high frequency than when they fire at low frequency. This hypothesis is supported by the fact that the spatial frequency-tuning curve is flatter in wildtype than in CKAMP44^−/−^ mice (**Fig. 4F**), which results from the damping effect exerted by CKAMP44 in relay neurons for high frequency retinal inputs (at low spatial frequencies). In contrast to what one might assume from the rather flat average tuning curve, dLGN RCs have preferred spatial frequencies as evidenced by the inspection of tuning curves of individual units (see 2 example tuning curves with different preferred spatial frequencies in *Extended data Fig. 5F*). To better compare the influence of CKAMP44 on the steepness of spatial frequency tuning, we generated tuning curves after sorting the response amplitudes of each unit from lowest (indicated as stimulus 1) to highest (indicated as stimulus 5). Response amplitudes were higher for all stimuli (i.e. form least preferred to most preferred) in CKAMP44^−/−^ than in wildtype mice for all the 5 stimuli (**Fig. 4G**). The linear regression on the sorted spatial frequency tuning curves for each unit showed that deletion of CKAMP44 increases the slope of the response amplitudes, suggesting that CKAMP44 exerts a divisive (and not a subtractive) gain control on signal transmission in the dLGN, which can be explained by the increased low-pass filter properties of retinogeniculate synapses in the presence of CKAMP44. Inspection of the distribution of stimulus-evoked firing rates suggested that presence of CKAMP44 reduces the firing of the most responsive cells in particular. The distribution of stimulus-evoked firing rates was non-Gaussian with a long tail towards high firing rates in both genotypes (*Extended data Fig. 5D-E*). However, this long tail was more pronounced in CKAMP44^−/−^ than in wildtype mice as indicated by the significantly increased interquartile ranges. This is further evidence for a decrease in low-pass filter properties of retinogeniculate synapses in CKAMP44-lacking mice. The lower number of highly responsive units in wildtype than in CKAMP44^−/−^ mice suggests that one function of AMPAR desensitization in retinogeniculate synapses is to avoid overactivation or saturation of dLGN RCs activity.

### CKAMP44 does not affect orientation and direction selectivity of RCs

Motion direction detection is a major feature of information processing in the visual pathway^24–28^. RCs of the dLGN responded with different degrees of direction- and/or orientation-selectivity to the presentation of drifting gratings. To analyze orientation or direction preference, we quantified the response amplitudes to the different directions for each unit at the preferred spatial frequency. To compare the response amplitudes to the different directions, we shifted the tuning curves of each unit such that the highest response is shown at direction 3. CKAMP44 deletion increased the mean firing rate for all directions (*Extended data Fig. 5J*). We then calculated the orientation and direction selectivity indices (DSI and OSI) for each unit at their preferred spatial frequency. DSI and OSI distributions were not different between genotypes (*Extended data Fig. 5G-H*). This may not be too surprising as this information is relayed from the retina to the visual cortex without major transformation by dLGN RCs^29^ and is consistent with our previous observation that CKAMP44 has no obvious influence on retina function^16^.

### CKAMP44 affects contrast tuning of dLGN RCs by decreasing input and response gain

The contrast gain of dLGN RCs is lower than that of RGCs which results from a transmission rate that interestingly decreases with increasing contrast (i.e. with the frequency of RGC activity^30^). Our simulations of RC activity (**Fig. 3J**) suggested that desensitization of AMPARs is responsible for this transformation of the contrast response curve. To test this assumption, we presented 5Hz full-screen ON-OFF stimuli with different levels of contrast. The responses of units of both genotypes are temporally locked to the stimulus and response amplitude increased with the contrast of the stimulus with an apparent higher peak frequency of units in CKAMP44^−/−^ than in wildtype mice (**Fig. 5A**). Deletion of CKAMP44 significantly increased the mean response amplitude at all the contrasts (**Fig. 5B**). In addition, a fitting with hyperbolic-ratio function (**Fig. 5C**) revealed that CKAMP44 deletion increases R_max_ by ≈54% and reduces C50 by 33%. Remarkably, the difference in R_max_ and C50 between genotypes was in a similar range to the values of the fitting of simulated input/output curves of RC firing (**Fig. 3J**). This suggests that CKAMP44 reduces input and response gain of the dLGN by increasing desensitization of AMPARs in retinogeniculate synapses.

## Discussion

In this paper, we investigated the role of CKAMP44 and AMPAR desensitization in information processing in the visual system. The generation of a RC model with retinogeniculate synapses containing AMPARs with different desensitization-kinetics allowed us to investigate important determinants of STP in the dLGN. The main finding of the simulations that AMPAR desensitization influences input and response gain of RCs was corroborated by *in vivo* recordings of dLGN neuron activity of wildtype and CKAMP44^−/−^ mice.

Visual information is not simply relayed through the visual thalamus. Instead, the likelihood of dLGN RCs activation depends on the history of RGC activity and it has been suggested that AMPAR desensitization affects signal transmission rate^30^. This assumption was based on *in vitro* data that showed pronounced STD of retinogeniculate synapses, which indeed depends on AMPAR desensitization. However, data from *in vivo* studies are contradictory with evidence of STD in retinogeniculate synapses in two studies^12,13^ and absence of relevant synaptic plasticity in another study^6^. Direct testing of the role of AMPAR desensitization in information processing is hampered by the difficulty to experimentally manipulate selectively AMPAR desensitization. Block of AMPAR desensitization by pharmacological tools is possible but affects also several other parameters of AMPAR and synapse function. For example, cyclothiazide, which efficiently blocks AMPAR desensitization, modulates also other gating kinetics^31^, increases vesicle release probability^32^ and induces robust epileptiform activity in rodents^33^. This precludes the use of this drug for the analysis of the role of desensitization in information processing *in vivo*. We therefore generated a computational model of RCs containing AMPARs with different levels of desensitization and analyzed activation of RCs in wildtype and CKAMP44^−/−^ mice, which differ in the amount of desensitization in retinogeniculate synapses^16^. The analyses showed that AMPAR desensitization indeed contributes substantially to STP in retinogeniculate synapses and to activation of RCs. The CKAMP44^−/−^ mouse constitutes a suitable model for the investigation of the role of AMPAR desensitization on signal transmission as the deletion of CKAMP44 reduces the magnitude of desensitization and recovery from desensitization substantially but does not alter deactivation and desensitization rates in dLGN neurons^16^. In addition, CKAMP44 does not affect active and passive membrane properties of dLGN RCs^16^. Finally, the influence of CKAMP44 on STD, membrane depolarization and neuron activation is also present when blocking NMDA or GABA receptors^16^, suggesting that increased activation by NMDA receptors or decreased inhibition by local interneurons do not account for the increased response amplitudes of RCs in CKAMP44^−/−^ mice. Deletion of CKAMP44 reduces synaptic AMPAR number by 43%^16^. Thus, the increase in RC activity would be even more pronounced if desensitization were reduced but synaptic strength unchanged. Indeed, this was observed in simulation experiments in which we reduced desensitization but not AMPARs number (data not shown). Thus, due to reduction in strength of retinogeniculate synapses in CKAMP44^−/−^ mice we in fact underestimate the relevance of AMPAR desensitization for signal transmission.

Vesicle release probability has been suggested to be considerably lower *in vivo* than *in vitro*^1^, which may well explain that STD is less pronounced *in vivo* than *in vitro*. In synapses without glutamate spillover, AMPAR desensitization modulates STP only when two vesicle releases occur in close temporal sequence in one release site. Consequently, the contribution of AMPAR desensitization to STP decreases with reduced vesicle release probability. In particular, the results of the study of Carandini and colleagues therefore questioned the relevance of STD and AMPAR desensitization for information processing in the dLGN^6^. Our simulations show that AMPAR desensitization affects STP not only under conditions with high vesicle release probability as observed *in vitro* but also in synapses with lower vesicle release probabilities that show little or no STD. Importantly, the simulations show that spillover of glutamate from active vesicle release sites to non-active release sites is essential for the influence of AMPAR desensitization on STP in synapses with low vesicle release probability.

The simulations showed that background activity (noise) influences the contribution of AMPAR desensitization to STP in retinogeniculate synapses. AMPAR recover fully from desensitization during prolonged time periods of synaptic inactivity. Spontaneous activity of retinogeniculate synapses is low in acute brain slices which may well explain why AMPAR desensitization shapes responses when the synapses is activated *in vitro* with several stimuli at high frequency. Differences in visual stimuli and/or background images may explain the different levels of STP *in vivo*. Thus, high background activity may result in constant high levels of AMPAR desensitization thereby reducing the contribution of further AMPAR desensitization during presentation of visual stimuli as previously suggested ^9^. Deletion of CKAMP44 decreases slightly the spontaneous firing rate of dLGN neurons. This may be explained by the fact that deletion of CKAMP44 reduces AMPAR number in retinogeniculate synapses by around 40%^16^, which would decrease excitation of RCs if the frequency of background (noise) activity is very low.

The visual thalamus functions as a spatio-temporal filter for visual information. Interestingly, the transmission rate decreases with the activity of retinal ganglion cells^30^. For example, although the firing rate of RGCs and dLGN RCs increase with contrast, fewer RGC spikes are transmitted at high contrast than at low contrast^30^. This decrease in response gain (response compression) has been suggested to result from saturation of RC activity and/or synaptic STD and AMPAR desensitization^30^. The lower R_max_ of the contrast tuning curves in wildtype than in CKAMP44^−/−^ mice suggests that indeed the decrease in response gain in the dLGN depends largely on AMPAR desensitization. CKAMP44 decreases not only R_max_ but also increases C50. This shows that AMPAR desensitization decreases not only response gain (reflected by the decrease in R_max_), but controls also contrast gain (reflected by the increase in C50). The magnitude in difference of R_max_ and C50 between genotypes was in a similar range to the difference of the same values of input-output curves between the two computational models. The only difference between the two models is the kinetics of AMPAR in retinogeniculate synapses. The data from the model therefore support that the difference in AMPAR desensitization in retinogeniculate synapses is the main factor explaining the difference in visual information processing between genotypes. The conclusion that AMPAR desensitization decreases response gain was confirmed by the analysis of drifting grating tuning curves. Also here, we observed that the differences in RC activity between wildtype and CKAMP44^−/−^ mice increases with the strength of RC activation. In other words, AMPAR desensitization decreases RC activity not in a subtractive but divisive way.

The likelihood that RCs are activated increases with the frequency of RGC activity with a steep increase in transmission rate with frequencies of more than 30 Hz^6–8,14,15^. RGC activity below 30Hz rarely elicits an action potential in RCs^8^. The increase in firing probability at high input frequencies can be explained with summation of EPSPs^9^. This frequency-dependent increase in transmission rate is very relevant for information processing in the dLGN as the second of two action potentials carries more visual information than the first when RGCs fire at high frequency^8,14,15^. The short memory of RCs therefore ensures that spikes with high information content but not noise are transmitted to RCs^14^. What role play synaptic STP and desensitization of AMPARs in controlling transmission rate? Recordings from RCs of CKAMP44^−/−^ mice showed that retinogeniculate synapses are presynaptically facilitating^16^. In wildtype neurons, STD therefore results from a balance of presynaptic facilitation and postsynaptic desensitization. Without the pronounced desensitization of AMPARs, short-term facilitation of retinogeniculate synapses together with summation of EPSPs may bear the risk of overactivation of RCs. Indeed, presence of CKAMP44 affects the distribution of amplitudes of RC responses by decreasing in particular the proportion of highly active RCs. In addition, CKAMP44 reduces the activation of RCs during presentation of non-preferred stimuli. This can be seen by 2Hz modulation of activity of RCs in wildtype mice during presentation of 2Hz drifting grating. Activity of RCs is close to background activity during presentation of the non-preferred signal of the grating. In contrast, activity remains higher throughout the presentation of drifting gratings in CKAMP44^−/−^ mice. AMPARs desensitization therefore seems to increase the encoding of temporal information of drifting gratings by decreasing the likelihood that action potentials of RGCs during presentation of the non-preferred signal (i.e. spikes with little information content) are transmitted to RCs. Finally, the control of the contrast gain exerted by CKAMP44 (increase in C50) suggests that AMPAR desensitization may enable RCs to better encode the different contrasts by preventing saturation of RC activation at the presentation of too low contrast images.

---

**Extended figure 1.**
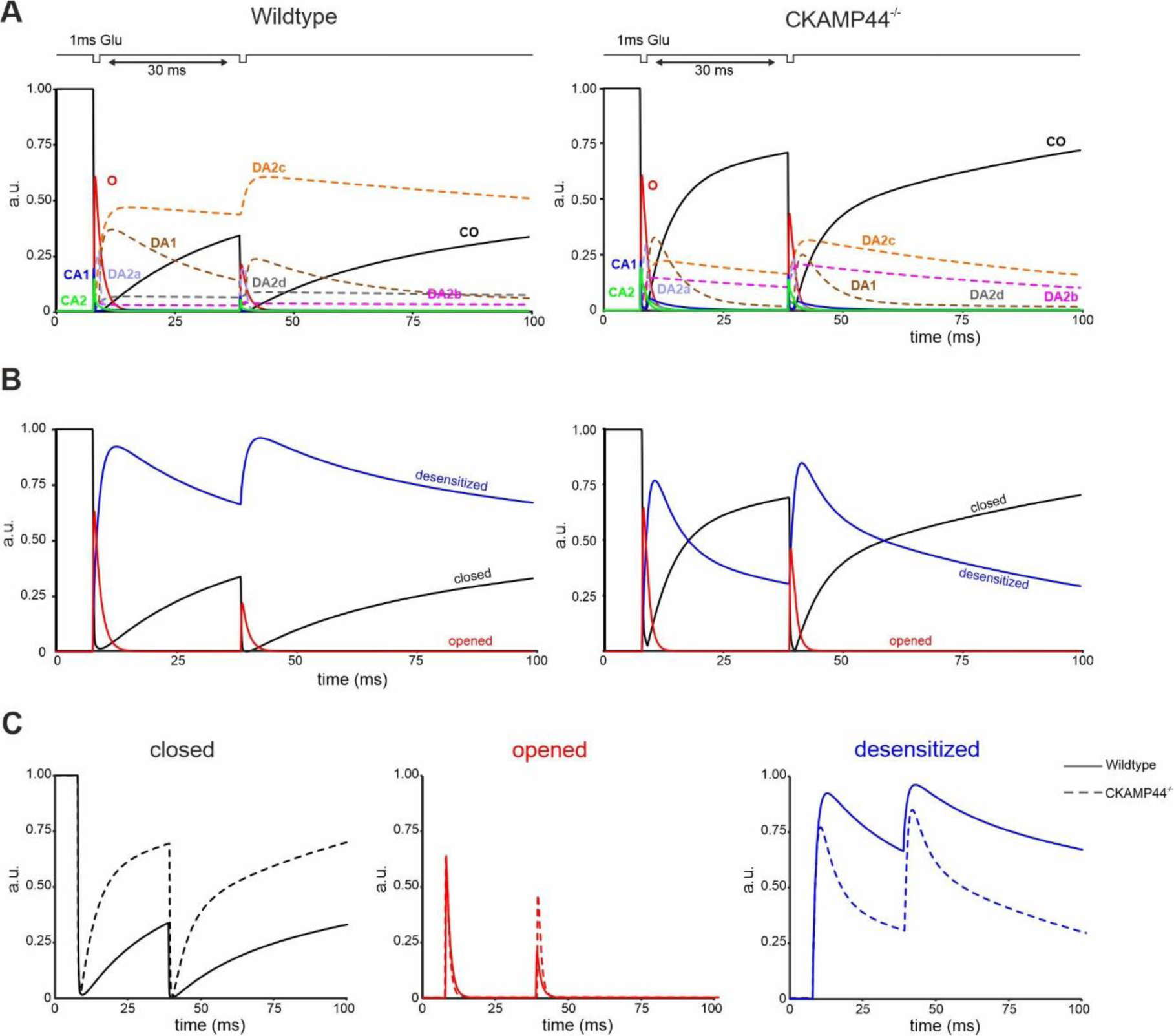
Time course of different states of AMPA receptors in response to two glutamate pulses. **A)** Time course of the nine states of wildtype (left) and CKAMP44^−/−^ (right) AMPA receptor model in response to two 1ms-pulses of 1mM glutamate with an inter-pulse interval of 30ms. At the second glutamate pulse, the CKAMP44^−/−^ AMPA receptor model displays higher open probability compared to the wildtype model. All desensitized states display, in absence of CKAMP44, a lower occupancy probability. In both genotypes, each state is represented with different colors. In addition, the open state and all closed states are indicated with continuous lines, whereas all desensitized states are represented with dashed lines. **B)** Time course of the open state (red), sum of all closed state (black) and sum of all desensitized states (blue) for wildtype (left) and CKAMP44^−/−^ (right) AMPA receptor model. **C)** Comparison wildtype vs. CKAMP44^−/−^ of the sum of all closed (right), open (middle), desensitized (right) states.

**Extended figure 2.**
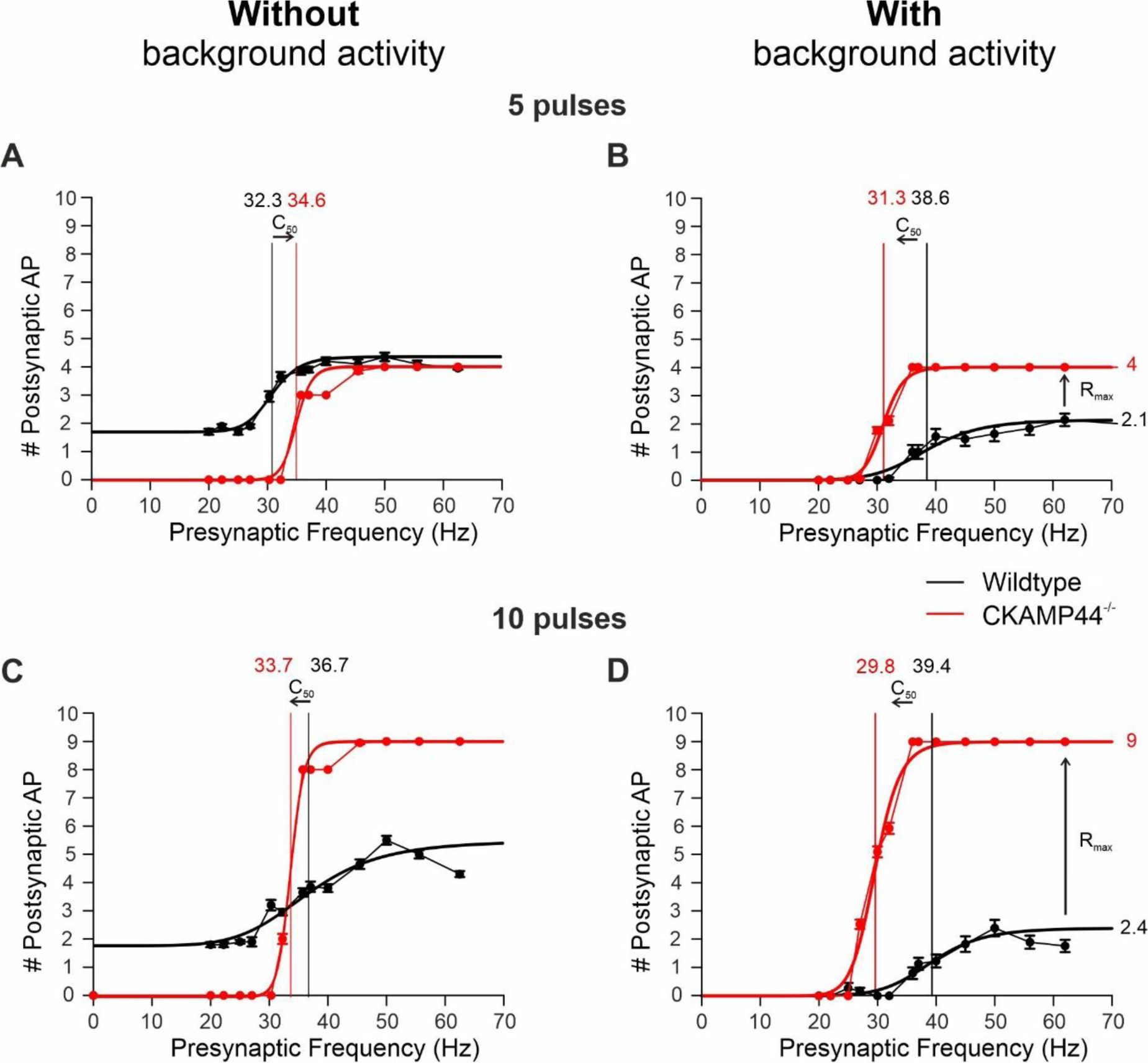
Model neuron activity with and without background activity. Simulations carried out similar as the ones shown in Figure 3 in absence **(A)** and in presence **(B)** of 2sec 10Hz background activity preceding activation of model neurons with five pulses at high frequency. **C)** and **D)** same as **A)** and **B),** respectively, with activation of model neurons with 10 pulses at high frequency.

**Extended figure 3.**
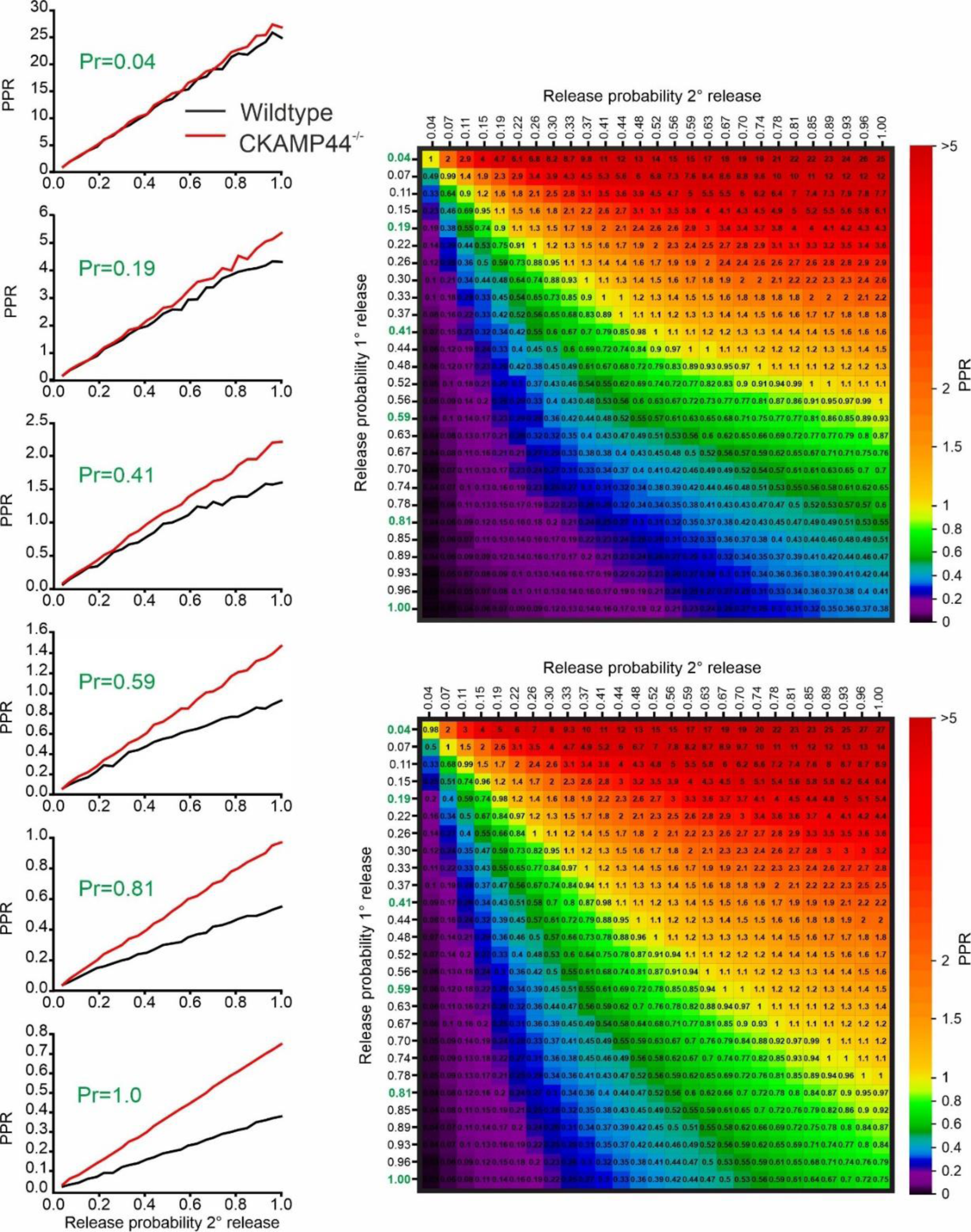
Dependence of the influence of desensitization on short-term plasticity on release probability. **(left)** Paired-pulse ratio plotted against release probability at second synapse activation for different release probability at first synapse activation. PPR is not influenced by different rates of recovery from desensitization (i.e. wildtype vs. CKAMP44^−/−^) if release probability at first synapse activation is low (top left, Pr=0,04 in green). The reduction in PPR with increasing release probability at first synapse activation (left, from top to bottom: Pr=0,04 to 1.0 in green) is more pronounced in the CKAMP44^−/−^ than in wild-type model. The difference in PPR between genotypes increases with release probability at first and second synapse activation. **(right)** Color map of the PPR for the wildtype (top right) and CKAMP44^−/−^ (right bottom) model. The release probabilities of the first synapse activation used for the plots on the left are marked in green on the left axis of the color maps. A faster recovery from desensitization (i.e. CKAMP44^−/−^ model) increases PPR almost uniformly for all release probabilities of first and second release.

**Extended figure 4.**
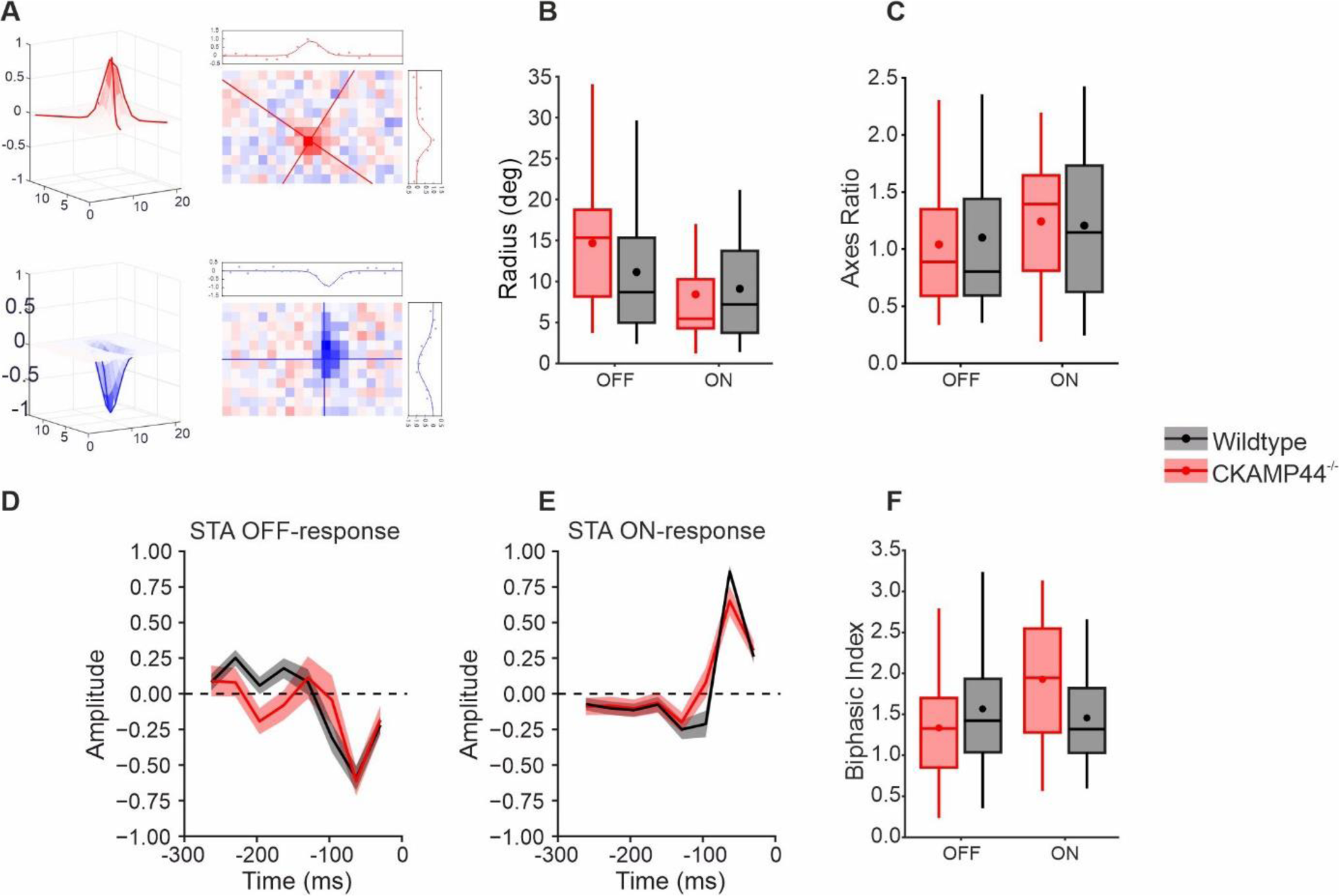
Genetic deletion of CKAMP44 does not influence receptive field shape of relay units. **A)** Spatial STA profile from an ON-responsive unit (top left) and an OFF-responsive unit (bottom left). The color map shows the intensity of the response to the white-noise stimulus. Each spatial STA profile was fitted with a 2-dimentional Gaussian (right). The lines in the color map indicate the x’ and y’ axis of the Gaussian function (see Methods). Plot on the top and on the right of spatial STA the represent the 2-d Gaussian section profile along the x- and y-axis respectively. **B and C)** Quantification of spatial STA measures**. B)** Average receptive field radius in ON- and OFF-responsive units from wildtype and CKAMP44^−/−^ mice (Wilcoxon test: *OFF -* W=328, p=0.16; *ON* – W=715, p=0.57, median ± IQR, dot = mean). **C**) Ratio between the horizontal and vertical radius in ON- and OFF-receptive fields of units from wildtype and CKAMP44^−/−^ mice (Wilcoxon test: *OFF -* W=411, p=0.90; *ON* – W=644, p=0.85, median ± IQR, dot = mean). **D + E)** Temporal STA response for OFF-responsive **(D)** and ON-responsive **(E)** units of wildtype and CKAMP44^−/−^ mice. Mean ± SEM. **F)** Biphasic index of the temporal STA of OFF- and ON-responsive units of wildtype and CKAMP44^−/−^ mice. The index was quantified as the ratio of maximum to minimum (ON-responsive units) or minimum to maximum (OFF-responsive units) response amplitude (Wilcoxon test: *OFF -* W=526, p=0.11; *ON* – W=534, p=0.17, median ± IQR, dot = mean). Total number of units: Wildtype n=89; CKAMP44^−/−^ n=48.

**Extended figure 5.**
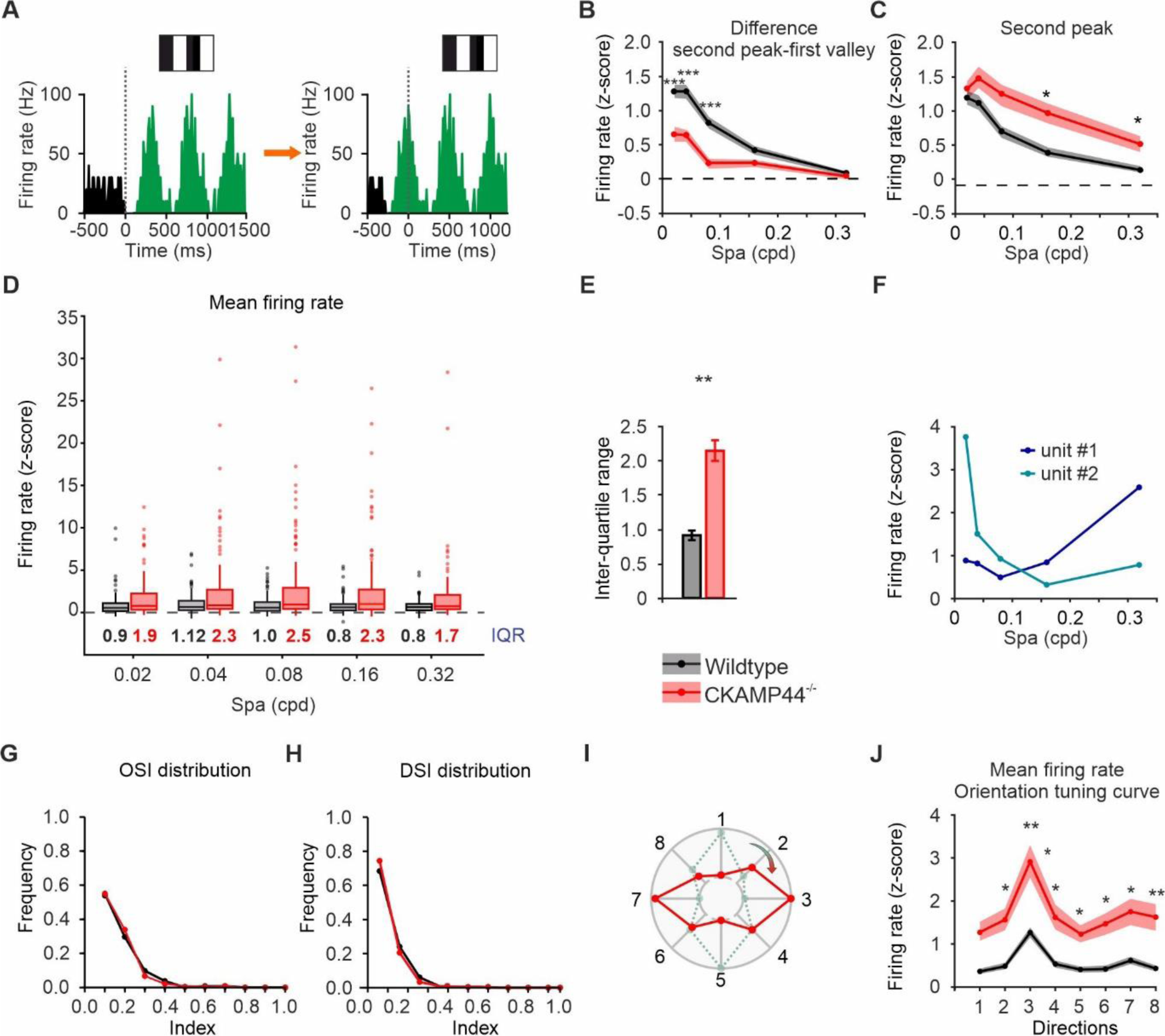
*In vivo* responses with OSI and DSI of relay cells to drifting gratings. **A)** Time series shift of ON- (top) and OFF- (bottom) responsive units to position the peak at 0ms. **B)** Difference between the second peak and the first valley amplitude plotted against spatial frequency (Wilcoxon test: 0.02cpd W=5779, p=0.000009; 0.04cpd W=5930, p=0.00003; 0.08cpd W=5729, p=0.000005; 0.16cpd W=7533, p=0.16, 0.32cpd W=8316, p=0.57, mean ± SEM). **C)** Amplitude of second peak of firing rates against spatial frequency (Wilcoxon test: 0.02cpd W=8848, p=0.54; 0.04cpd W=8845, p=0.58; 0.08cpd W=9485, p=0.1013;.16cpd W=9693, p=0.02, 0.32cpd W=9354, p=0.02, mean ± SEM). **D)** Boxplot of z-score mean firing rate at the best (preferred) direction of each unit. The values in the plot indicate the IQR of units from wildtype (black) and CKAMP44^−/−^ (red) mice. CKAMP44^−/−^ units display an increased IQR at all spas. **E)** Bar plot of IQRs (Wilcoxon test: W=0, p=0.008, mean ± SEM). **F)** Spatial frequency tuning curves of two different units recorded in the same mouse. Unit#1 shows the highest response at 0.32 cpd and unit#1 at 0.02 cpd. **G & H)** Normalized OSI and DSI distributions (two-sample Kolmogorov-Smirnov test: *OSI* – D=0.2 p=0.96, *DSI* – D=0.2 p=0.98). **I)** The tuning curve of this example unit (dashed green line) was shifted in order to display the highest response at direction 3 (continuous red line). **J)** Average orientation tuning curves of units from wildtype (black) and CKAMP44^−/−^ (red) mice (Wilcoxon test: *Direction 1* W=7548, p=0.102, *Direction 2* W=7127, p=0.02, *Direction 3* W=6773.5, p=0.004, *Direction 4* W=7167, p=0.02, *Direction 5* W=7287.5, p=.04, *Direction 6* W=7247, p=0.03, *Direction 7* W=7225, p=0.03, *Direction 8* W=6883.5, p=0.006, mean ± SEM). Number of units for all plots in the panel: Wildtype n=123, CKAMP44^−/−^ n=139.

## Materials and Methods

### Animals and ethical compliance

All animal experiments were performed in accordance with local Rheinland-Pfalz welfare regulation. All the animals used in these experiments belong to the C57BL/6N substrain. CKAMP44^−/−^ animals were generated as described previously^20^. In vivo recording of neuron activity was performed with 3-4 months old C57BL/6N mice (8 wildtype and 8 CKAMP44^−/−^). Animals were single housed and kept on a 12-h dark-light cycle. Recordings took place during the dark phase.

### Surgical procedure

Recording drives were lightweight 8-tetrodes (32-channel) headstages (Axona) and the tungsten tetrodes could be moved along the z-axis via screws with a resolution of 250 µm per turn. Tetrodes impedance was lowered to 300-400 kΩ by golden-plating tetrodes tips (gold chloride solution, SIGMA-ALDRICH), to reduce signal-to-noise ratio.

Animals were anesthetized with isoflurane (3% induction and 1.5-1.8% maintenance) and placed in a stereotaxic-apparatus. Body temperature was maintained at 37° via controlled heating pad. Eyes were covered with Bepanthen (Bayer, Germany) during the surgery. As soon as the absence of pedal and tail-pinch reflexes confirmed a deep state of anesthesia, the skin of the head was disinfected with antiseptic and then shaved. The skull was exposed and treated with Gel Etchant (37.5% Phosphoric acid, Kerr). Four screw holes were then drilled on the skull surface. Two of the screws were placed on the interparietal bone and then connected to a copper wire, serving as grounding and reference signal. The other two screws were fixed to the frontal lobe to increase stability to the implant. After drilling through skull, tetrodes were inserted to a depth of 2.6 mm using the following coordinates: 2.1 mm from sagittal suture and −2.1 mm from bregma. The microdrive and screws were fixed to the skull using dental cement. During the three post-operative days, intraperitoneal injection of Carprofen (5mg/kg a.c.) and Burprofen (0.1mg/kg) was given to the mouse two times per day. The animals were allowed to recovery for 1-2 weeks post-surgery before starting the experiments.

### Recording box and Visual Stimuli

After recovery time post-surgery, the mice were habituated to the head-fixation in the recording setup. The *in vivo* recordings were carried out in a customized recording cage (dimensions: 45×58×35 cm). A 24-inch monitor (XL2411Z, BenQ) was placed ≈25cm away from the head-fixed non-anaesthetized mouse with a viewing angle of ∼45° on left and right side, ∼37° above and below. Visual stimuli were presented on a gamma-corrected LED monitor (resolution 1920×1080) with a refresh rate of 60 Hz and were controlled by a custom-made software (Visual C++ and OpenGL).

To analyze receptive field features, a binary spatiotemporal white noise in a checkerboard layout was displayed on the monitor. Each square had a size of 96×96 pixels, and was updated randomly to black or white at 100% contrast at 30 Hz^34^. The evoked activity was recorded for ≈15 min.

Full-field square-wave drifting gratings (100% contrast, temporal frequency of 2Hz) were presented for 1.5s at eight fixed evenly spaced motion direction and five spatial frequencies (0.02, 0.04, 0.08, 0.16 and 0.32 cpd), summing up to 40 different stimuli, similar as described previously ^24,35–37^. Before and between presentation of moving gratings, the monitor was set for 3 s to full-field gray (50% contrast) to estimate background activity and to allow recovery from stimulus presentation. Each of the 40 different drifting grating stimuli was presented 10 times in a random order.

To analyze sensitivity and gain control properties of relay cells, stimuli with different contrasts were presented. The stimulus consisted of 2s background (full-field gray, 50% contrast) followed by 1s where the luminance alternates at the frequency of 10 Hz between 60-40 %, 70-30 %, 80-20 %, 90-10 %, or 100-0 % of the maximal monitor luminance. Considering only the variation between the two levels of contrast, the five contrast steps analyzed in this study are 20%, 40%, 60%, 80%, and 100%. The stimulus was repeated 10 times.

### Data Acquisition

Tetrodes were lowered into the dLGN by ≈100-150 µm/per day over multiple days in order to increase the number of recorded neurons. Each recording session was performed 24h after lowering the position of tetrodes. The signal was amplified and digitized at 20 kHz (RHD2000-Series, Amplifier Evaluation System, Intan Technologies, with analog bandwidth 0.09Hz-7.6Hz). The correct position of tetrodes in the dLGN was identified by a strong increase in activity of recorded units in response to full-field ON-OFF stimuli. In addition, the correct position of the tetrodes was confirmed by histological analysis postmortem.

Spike detection and sorting were performed using Klusta^38^ (https://klusta.readthedocs.io/en/latest/). Klusta takes the binary raw data file obtained from RHD2000 system recordings and performs spike detection and sorting after band-pass filtering the raw signal (0.8-3 kHz). The threshold for spike detection was set to 5-fold of the SD of each recorded signal. Spikes waveforms were extracted from the four wires of each tetrode and then processed in the automatic clustering algorithm to group the spikes into putative neuronal sources. Results from the automatic clustering were then manually refined by using a graphical user interface (Phy). Clusters not violating the 2ms refractory period were considered as single units and kept for further analysis. Finally, a peristimulus time histogram with 10ms bin was constructed for all the single units obtained after manual refinement.

### Data Analysis

All data were analyzed by using custom scripts written in R and in MATLAB. Spike-triggered analysis (STA) of visual stimuli is the average stimulus that precedes a spike and it was used to estimate receptive field of RCs ^39,40^. Several cells exhibit receptive fields, whose spatiotemporal structure can be expressed as the product of a temporal and a spatial function, which can be treated as independent variables^24,34,40–42^. Cells that did not show a spatial STA component with amplitude higher than the background were not included in RF analysis. The spatial STA was fitted by a 2-dimensional Gaussian distribution with independent standard deviations (*σ*_*x*_, *σ*_*y*_) which were then used to quantify the receptive field size, shape and polarity. The temporal STA was then used to evaluate the biphasic nature of the response by calculating the ratio between the minimum and maximum stimulus amplitude of both ON- and OFF-cells.

For drifting gratings and contrast stimuli, after construction of peristimulus histograms for each unit and each stimulus condition, the number of spikes in each bin were divided by the bin size and averaged across the number of trials, thus obtaining firing rates. As mentioned above, automated spike sorting was followed by user manual curation, from which ≈500 units were obtained from wildtype and CKAMP44^−/−^ mice, respectively. Among the detected units, we selected for further analyses those neurons that were responsive to visual stimuli. External stimuli can induce changes in firing rate with respect to the mean spontaneous activity as well as responses that are modulated by stimulus frequency. We selected units showing a clear response to the visual stimulation, applying the following criterion. A Fourier analysis on the firing rate trace was performed in the 1000ms interval after stimulus onset. The F0 component informs about changes in the mean firing rate. In addition, the firing rate of relay neurons can modulate their response at the first harmonic (F1 component). Therefore, responsive units often show an F1 component that is larger than other frequency components. To identify units that respond to the visual stimulus, we applied a z-score analysis approach on the first 15 Fourier components.

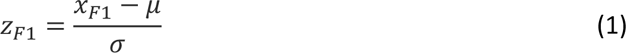

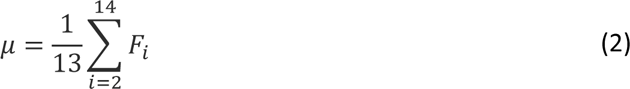

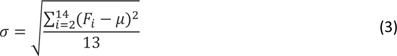

Eq. (2) represent the average *μ* and the standard deviation *σ*, respectively, of all 15 Fourier components without the amplitudes of F0 and F1 components. Therefore *z*_*F*1_ informs about the magnitude of firing rate modulation to the visual stimulus. Finally, we selected responsive units based on thresholds (identical for both genotypes) for *z*_*F*1_ and *z*_*F*0_. The latter indicates the z-score of the F0 component with respect to the average of 50 F0 components calculated over 50 sliding windows of 1000ms length using 10ms increment starting from −1500ms (i.e. the average of the 50 F0s is considered the background F0). This criterion was applied for responses to contrast stimuli and moving grating stimuli independently. A unit was considered responsive to moving gratings if its activity upon visual stimulation met or exceeded a threshold value (independent for *z*_*F*1_ and *z*_*F*0_) at least in one of the 40 possible combinations (8 directions and 5 spatial frequencies). Based on these criteria, the number of units that were responsive to moving gratings were 123 wildtype and 139 CKAMP44^−/−^. Likewise, a unit was considered responsive to contrast stimuli if responsive at least to one of the five contrasts used. Based on these criteria, 83 wildtype and 44 CKAMP44^−/−^ units were responsive to contrast stimuli.

### Tuning curves

To characterize the unit activity evoked by the different stimuli, we generated tuning curves by plotting the z-score of firing rate calculated as a function of stimulus parameters. In both moving gratings and contrast stimuli, the z-score was calculated by Eq. (4):

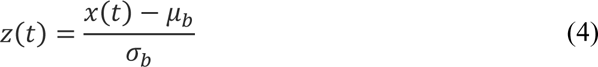

Where *x*(*t*) is the response amplitude of a neuron in the 1500ms after stimulus onset, and *μ*_*a*_ and *b*_*a*_ are the mean and standard deviation, respectively, of the baseline activity calculated from time −1000 to 0 ms. When generating moving gratings tuning curves, the average z-score of the firing rate was calculated for each combination of directions and spatial frequency (= 40 different z-scores per unit). Spatial frequency-tuning curve was built by plotting the average z-score firing rate against spatial frequency at the direction where the response was the highest (best direction). The orientation tuning curve was obtained by plotting the z-score firing rate against the eight directions at the spatial frequency where the response was the highest (best spatial frequency). The preferred direction of motion can differ among neurons: for this reason, the orientation tuning curves per single unit have been shifted in order to have the highest response (across the eight directions) at direction number 3 (*Extended Data Fig.5-I*).

Orientation Selectivity Index (OSI) and Direction Selectivity Index (DSI) were calculated using a standard metric based on the circular variance as follows:

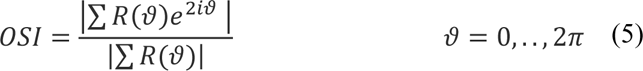

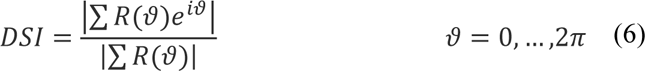

In both equations, *R*(*ϑ*) represents the single unit mean response at direction *ϑ*. The absolute amplitudes of these values return OSI and DSI.

To obtain contrast tuning curve, the z-score firing rate was plotted against the five different contrasts used, thus obtaining contrast-tuning curves. In order to analyze gain control properties of relay neurons, the hyperbolic ratio function was fitted to the contrast-tuning curves.

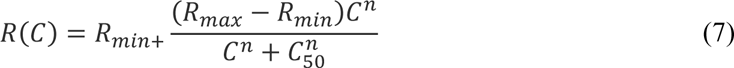

In Eq. (7), *R*_*max*_ and *R*_*min*_ represents the maximal and the minimal response among all the contrasts presented respectively, *n* indicates the sensitivity and *C*_50_represent the semi-saturation contrast value (50% of *R*_*max*_)^43^.

### Statistics and data presentation

Statistical analysis was performed using R-Studio software. To evaluate whether data are normally distributed, Shapiro-Wilk test was used. Data normally distributed were compared using student t-test. Data not normally distributed were statistically compared using Mann-Whitney test. Data are shown as mean and standard error of the mean or as median and interquartile range.

### Computational model

Simulation were carried out in NEURON simulation environment for modelling of individual neurons as well as networks of neurons^44^. The main goal of the computational modelling implemented in this study was to better understand the relevance of mechanisms (AMPA receptors desensitization, spillover of glutamate, vesicle release probability) involved in the information processing at the retinogeniculate synapses. Therefore, the following model was implemented: upon activation a single RGCs releases glutamate molecules that diffuse in the synaptic cleft and bind AMPARs located in the membrane of a RCs. The total increase of AMPARs-current induces the depolarization of RCs membrane, leading to action potentials (APs) if the threshold is reached.

A simplified multi-compartmental representation of a RCs was adapted from an established model from Destexhe and colleagues: a realistic morphology of RCs was collapsed into three compartments by applying the axial resistance conservation method^45^. Moreover, in order to compensate for potential incorrect input resistance, conductance and capacitance of each compartment (except for the soma) were multiplied by a dendritic correction factor *C*_*d*_ of 7.95, as done in^45^. To the three-compartment simplified model of the relay cell, an additional dendritic arborization composed of 24 compartments was connected. The structure was built in order to obtain a specific two-dimensional distribution of AMPAR clusters in the post-synaptic terminal of the retinogeniculate synapse. This structure was based on parameters obtained from published results of electron microscopy 3D reconstructions of retinogeniculate synapses^5^. The scheme in **Fig. 2A- and B**-left shows the distribution of the 27 AMPARs clusters used in this model. AMPARs arrangement reflects experimental evidence from Budisantoso and colleagues^5^. In their publication they showed that the number of neighbor synapses within 700nm and 1400nm is respectively 1 and 6^46^. Vesicle releases sites of axon terminals of the retinal ganglion cell were located opposite of each AMPA receptor cluster with a synaptic cleft of 20 nm.

In this study, we considered only the axonal terminal of the retinal ganglion cell, with a total area of 113.15 µm^2^. Importantly, passive and active properties of the retinal ganglion cell, such as leak channel density and voltage gated channels distribution, were set so that the cell fires action potentials upon stimulation. Leak channels had a reversal potential *E*_*L*_ = −70 *mV* and a peak conductance of *g*_*L*_ = 0.0003 *S*/*cm*^2^. The axial resistance *R*_*a*_ was set to 173 *Ωcm*. Sodium and potassium reversal potentials were set to *E*_*Na*_ = 55 *mV* and *E*_*K*_ = −90 *mV* and their peak conductance were set to *g*_*Na*_ = 0.01 *S*/*cm*^2^ and *g*_*K*_ = 0.003 *S*/*cm*^2^, respectively, which is in line with previous work (i.e.^47^).

The simplified model of RCs has a total area of 6029.5 µm^2^, which is similar to the previous established model ^45^. Passive properties of RCs were obtained from fitting the experimental input resistance. Fixed the reversal potential of leak channels *E*_*L*_ = −70 *mV*, the peak conductance *g*_*L*_ of the RCs model was set to 0.00003 *S*/*cm*^2^ such that the slope of the IV curve obtained by injecting hyperpolarizing currents would fit the experimental results from Chen and colleagues^16^ (not shown). Sodium and potassium reversal potentials were set to *E*_*Na*_ = 55 *mV* and *E*_*K*_ = −90 *mV* respectively. Peak conductance were *g*_*Na*_ = 0.27 *S*/*cm*^2^ and *g*_*K*_ = 0.0215 *S*/*cm*^2^, which are in the range of previous works^45,47,48^. Using these parameters, response amplitude to hyperpolarizing currents of the relay cell model was in a similar range as observed in *in vitro* experiments^16^.

### Glutamate release and AMPA receptor model

In order to simulate ultra-fast application of glutamate, a simple release process has been implemented, where inter-pulse interval, duration of each pulse and amount of glutamate released could be regulated. We set the glutamate concentration to 1mM and considered five different interpulse intervals (30ms, 100ms, 300ms, 1000ms and 3000ms) of 1ms long pulse. A single stimulation with 100ms long pulse was also considered. The experimental data were fit by using the NEURON function *Multiple Run Fitter*. To simulate input transmission in the retinogeniculate synaptic model presented in this study, we implemented a well-established release mechanism fully described by Destexhe and colleagues^47^. From experimental results, it has been observed that the release probability a retinogeniculate synapses varies between 0.3 and 0.7^1^. In the model presented in this study, the release probabilities at the different stimuli was set consistently with experimental data from our laboratory suggesting that release probability increases slightly when retinogeniculate synapses are activated several times in a short time period^16^. The initial release probability (Pr) was set to ≈ 0.37. Glutamate diffusion in the synaptic cleft of retinogeniculate synapses was simulated by using rxd-module in NEURON^49,50^. Tortuosity and fractional volume were set to 1. Diffusion coefficient D in the synaptic cleft was set to 0.46 µm^2^/ms, according to recently published experimental results^18^.

The process described in the previous paragraph was used to fit the experimental AMPAR-mediated currents evoked by ultra-fast application of glutamate onto nucleated patches of relay cells from Chen and colleagues^16^. The experimental response to a single pulse was modeled using the Eq. (8), where all constants were set in order to fit the average amplitude and decay time of experimental waveforms^16^.

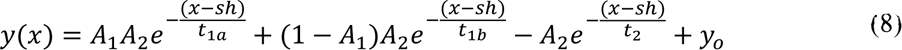

We adopted a well-known nine-state AMPA receptor kinetics model (Fig. 3A)^17,46^. Each transition between two states can be described by kinetic equations following Markovian formalism. The equations set was implemented in NEURON using NMDOL language. Therefore, the optimization process described above was run in order to find a set of rate constants such that the AMPA receptor currents from the model fit the average experimental results from^16^.

## Acknowledgements

This work was funded by the German Research Foundation (DFG) grant within the Collaborative Research Center (SFB) 1080 “Molecular and Cellular Mechanisms of Neural Homoeostasis” to JvE. The authors want to thank Barbara Biesalski and Chiara Ardigò for excellent technical assistance.

## Author Contributions

S.R. developed the computational model, S.R. performed in vivo recordings, S.R. analyzed in vivo experimental data, T.G. provided visual stimulation protocols, S.R. and J.vE. conceived model and experiments and wrote the manuscript with inputs from T.G.

## References

1 Borst, J. G. The low synaptic release probability in vivo. Trends Neurosci 33, 259–266, doi:10.1016/j.tins.2010.03.003 (2010).

2 von Engelhardt, J. Role of AMPA receptor desensitization in short term depression - lessons from retinogeniculate synapses. J Physiol, doi:10.1113/JP280878 (2021).

3 Chen, C., Blitz, D. M. & Regehr, W. G. Contributions of receptor desensitization and saturation to plasticity at the retinogeniculate synapse. Neuron 33, 779–788, doi:10.1016/s0896-6273(02)00611-6 (2002).

4 Chen, C. & Regehr, W. G. Developmental remodeling of the retinogeniculate synapse. Neuron 28, 955–966, doi:10.1016/s0896-6273(00)00166-5 (2000).

5 Budisantoso, T., Matsui, K., Kamasawa, N., Fukazawa, Y. & Shigemoto, R. Mechanisms underlying signal filtering at a multisynapse contact. The Journal of neuroscience : the official journal of the Society for Neuroscience 32, 2357–2376, doi:10.1523/JNEUROSCI.5243-11.2012 (2012).

6 Carandini, M., Horton, J. C. & Sincich, L. C. Thalamic filtering of retinal spike trains by postsynaptic summation. Journal of vision 7, 20 21–11, doi:10.1167/7.14.20 (2007).

7 Mastronarde, D. N. Two classes of single-input X-cells in cat lateral geniculate nucleus. II. Retinal inputs and the generation of receptive-field properties. J Neurophysiol 57, 381–413, doi:10.1152/jn.1987.57.2.381 (1987).

8 Sincich, L. C., Adams, D. L., Economides, J. R. & Horton, J. C. Transmission of spike trains at the retinogeniculate synapse. The Journal of neuroscience : the official journal of the Society for Neuroscience 27, 2683–2692, doi:10.1523/JNEUROSCI.5077-06.2007 (2007).

9 Sincich, L. C., Horton, J. C. & Sharpee, T. O. Preserving information in neural transmission. The Journal of neuroscience : the official journal of the Society for Neuroscience 29, 6207–6216, doi:10.1523/JNEUROSCI.3701-08.2009 (2009).

10 Kielland, A. & Heggelund, P. AMPA and NMDA currents show different short-term depression in the dorsal lateral geniculate nucleus of the rat. J Physiol 542, 99–106, doi:10.1113/jphysiol.2002.019240 (2002).

11 Kielland, A. et al. Activity patterns govern synapse-specific AMPA receptor trafficking between deliverable and synaptic pools. Neuron 62, 84–101, doi:10.1016/j.neuron.2009.03.001 (2009).

12 Eysel, U. T. Quantitative studies of intracellular postsynaptic potentials in the lateral geniculate nucleus of the cat with respect to optic tract stimulus response latencies. Exp Brain Res 25, 469–486, doi:10.1007/BF00239782 (1976).

13 Weyand, T. G. Retinogeniculate transmission in wakefulness. J Neurophysiol 98, 769–785, doi:10.1152/jn.00929.2006 (2007).

14 Rathbun, D. L., Warland, D. K. & Usrey, W. M. Spike timing and information transmission at retinogeniculate synapses. The Journal of neuroscience : the official journal of the Society for Neuroscience 30, 13558–13566, doi:10.1523/JNEUROSCI.0909-10.2010 (2010).

15 Usrey, W. M., Reppas, J. B. & Reid, R. C. Paired-spike interactions and synaptic efficacy of retinal inputs to the thalamus. Nature 395, 384–387, doi:10.1038/26487 (1998).

16 Chen, X., Aslam, M., Gollisch, T., Allen, K. & von Engelhardt, J. CKAMP44 modulates integration of visual inputs in the lateral geniculate nucleus. Nature communications 9, 261, doi:10.1038/s41467-017-02415-1 (2018).

17 Hausser, M. & Roth, A. Dendritic and somatic glutamate receptor channels in rat cerebellar Purkinje cells. J Physiol 501 (Pt 1), 77–95, doi:10.1111/j.1469-7793.1997.077bo.x (1997).

18 Zheng, K. et al. Nanoscale diffusion in the synaptic cleft and beyond measured with time-resolved fluorescence anisotropy imaging. Sci Rep 7, 42022, doi:10.1038/srep42022 (2017).

19 Khodosevich, K. et al. Coexpressed auxiliary subunits exhibit distinct modulatory profiles on AMPA receptor function. Neuron 83, 601–615, doi:10.1016/j.neuron.2014.07.004 (2014).

20 von Engelhardt, J. et al. CKAMP44: a brain-specific protein attenuating short-term synaptic plasticity in the dentate gyrus. Science 327, 1518–1522, doi:10.1126/science.1184178 (2010).

21 Silver, R. A. Neuronal arithmetic. Nat Rev Neurosci 11, 474–489, doi:10.1038/nrn2864 (2010).

22 Tang, J., Ardila Jimenez, S. C., Chakraborty, S. & Schultz, S. R. Visual Receptive Field Properties of Neurons in the Mouse Lateral Geniculate Nucleus. PloS one 11, e0146017, doi:10.1371/journal.pone.0146017 (2016).

23 Cheng, H., Chino, Y. M., Smith, E. L., 3rd, Hamamoto, J. & Yoshida, K. Transfer characteristics of lateral geniculate nucleus X neurons in the cat: effects of spatial frequency and contrast. J Neurophysiol 74, 2548–2557, doi:10.1152/jn.1995.74.6.2548 (1995).

24 Piscopo, D. M., El-Danaf, R. N., Huberman, A. D. & Niell, C. M. Diverse visual features encoded in mouse lateral geniculate nucleus. The Journal of neuroscience : the official journal of the Society for Neuroscience 33, 4642–4656, doi:10.1523/JNEUROSCI.5187-12.2013 (2013).

25 Durand, S. et al. A Comparison of Visual Response Properties in the Lateral Geniculate Nucleus and Primary Visual Cortex of Awake and Anesthetized Mice. The Journal of neuroscience : the official journal of the Society for Neuroscience 36, 12144–12156, doi:10.1523/JNEUROSCI.1741-16.2016 (2016).

26 Baden, T. et al. The functional diversity of retinal ganglion cells in the mouse. Nature 529, 345–350, doi:10.1038/nature16468 (2016).

27 Cruz-Martin, A. et al. A dedicated circuit links direction-selective retinal ganglion cells to the primary visual cortex. Nature 507, 358–361, doi:10.1038/nature12989 (2014).

28 Sclar, G. & Freeman, R. D. Orientation selectivity in the cat’s striate cortex is invariant with stimulus contrast. Exp Brain Res 46, 457–461, doi:10.1007/BF00238641 (1982).

29 Scholl, B., Tan, A. Y., Corey, J. & Priebe, N. J. Emergence of orientation selectivity in the Mammalian visual pathway. The Journal of neuroscience : the official journal of the Society for Neuroscience 33, 10616–10624, doi:10.1523/JNEUROSCI.0404-13.2013 (2013).

30 Kaplan, E., Purpura, K. & Shapley, R. M. Contrast affects the transmission of visual information through the mammalian lateral geniculate nucleus. J Physiol 391, 267–288, doi:10.1113/jphysiol.1987.sp016737 (1987).

31 Partin, K. M., Fleck, M. W. & Mayer, M. L. AMPA receptor flip/flop mutants affecting deactivation, desensitization, and modulation by cyclothiazide, aniracetam, and thiocyanate. The Journal of neuroscience : the official journal of the Society for Neuroscience 16, 6634–6647, doi:10.1523/JNEUROSCI.16-21-06634.1996 (1996).

32 Diamond, J. S. & Jahr, C. E. Asynchronous release of synaptic vesicles determines the time course of the AMPA receptor-mediated EPSC. Neuron 15, 1097–1107, doi:10.1016/0896-6273(95)90098-5 (1995).

33 Qi, J., Wang, Y., Jiang, M., Warren, P. & Chen, G. Cyclothiazide induces robust epileptiform activity in rat hippocampal neurons both in vitro and in vivo. J Physiol 571, 605–618, doi:10.1113/jphysiol.2005.103812 (2006).

34 Schreyer, H. M. & Gollisch, T. Nonlinear spatial integration in retinal bipolar cells shapes the encoding of artificial and natural stimuli. Neuron 109, 1692–1706 e1698, doi:10.1016/j.neuron.2021.03.015 (2021).

35 Liu, J. K. et al. Inference of neuronal functional circuitry with spike-triggered non-negative matrix factorization. Nature communications 8, 149, doi:10.1038/s41467-017-00156-9 (2017).

36 Khani, M. H. & Gollisch, T. Linear and nonlinear chromatic integration in the mouse retina. Nature communications 12, 1900, doi:10.1038/s41467-021-22042-1 (2021).

37 Niell, C. M. & Stryker, M. P. Highly selective receptive fields in mouse visual cortex. The Journal of neuroscience : the official journal of the Society for Neuroscience 28, 7520–7536, doi:10.1523/JNEUROSCI.0623-08.2008 (2008).

38 Rossant, C. et al. Spike sorting for large, dense electrode arrays. Nat Neurosci 19, 634–641, doi:10.1038/nn.4268 (2016).

39 Schwartz, O., Pillow, J. W., Rust, N. C. & Simoncelli, E. P. Spike-triggered neural characterization. Journal of vision 6, 484–507, doi:10.1167/6.4.13 (2006).

40 Chichilnisky, E. J. A simple white noise analysis of neuronal light responses. Network 12, 199–213 (2001).

41 Ravi, S., Ahn, D., Greschner, M., Chichilnisky, E. J. & Field, G. D. Pathway-Specific Asymmetries between ON and OFF Visual Signals. The Journal of neuroscience : the official journal of the Society for Neuroscience 38, 9728–9740, doi:10.1523/JNEUROSCI.2008-18.2018 (2018).

42 Wolfe, J. & Palmer, L. A. Temporal diversity in the lateral geniculate nucleus of cat. Vis Neurosci 15, 653–675, doi:10.1017/s0952523898154068 (1998).

43 Albrecht, D. G. & Hamilton, D. B. Striate cortex of monkey and cat: contrast response function. J Neurophysiol 48, 217–237, doi:10.1152/jn.1982.48.1.217 (1982).

44 Hines, M. L. & Carnevale, N. T. The NEURON simulation environment. Neural Comput 9, 1179–1209, doi:10.1162/neco.1997.9.6.1179 (1997).

45 Destexhe, A., Neubig, M., Ulrich, D. & Huguenard, J. Dendritic low-threshold calcium currents in thalamic relay cells. The Journal of neuroscience : the official journal of the Society for Neuroscience 18, 3574–3588 (1998).

46 Budisantoso, T. et al. Evaluation of glutamate concentration transient in the synaptic cleft of the rat calyx of Held. J Physiol 591, 219–239, doi:10.1113/jphysiol.2012.241398 (2013).

47 Destexhe, A., Mainen, Z. F. & Sejnowski, T. J. Synthesis of models for excitable membranes, synaptic transmission and neuromodulation using a common kinetic formalism. Journal of computational neuroscience 1, 195–230, doi:10.1007/BF00961734 (1994).

48 Destexhe, A., Bal, T., McCormick, D. A. & Sejnowski, T. J. Ionic mechanisms underlying synchronized oscillations and propagating waves in a model of ferret thalamic slices. J Neurophysiol 76, 2049–2070, doi:10.1152/jn.1996.76.3.2049 (1996).

49 McDougal, R. A., Hines, M. L. & Lytton, W. W. Reaction-diffusion in the NEURON simulator. Frontiers in neuroinformatics 7, 28, doi:10.3389/fninf.2013.00028 (2013).

50 Newton, A. J. H., McDougal, R. A., Hines, M. L. & Lytton, W. W. Using NEURON for Reaction-Diffusion Modeling of Extracellular Dynamics. Frontiers in neuroinformatics 12, 41, doi:10.3389/fninf.2018.00041 (2018).

